# Voltage-driven polyelectrolyte complexation inside a nanopore

**DOI:** 10.1101/2021.06.21.449295

**Authors:** Prabhat Tripathi, Byoung-jin Jeon, Murugappan Muthukumar

## Abstract

We have investigated how a pair of oppositely charged macromolecules can be driven by an electric field to form a polyelectrolyte complex inside a nanopore. To observe and isolate an individual complex pair, a model protein nanopore, embedded in artificial phospholipid membrane, allowing compartmentalization (*cis*/*trans*) is employed. A polyanion in the *cis* and a polycation in the *trans* compartments are subjected to electrophoretic capture by the pore. We find that the measured ionic current across the pore has a distinguishable signature of complex formation, which is different from the signature of the passage of individual molecules through the pore. The ionic current signature allows us to detect the interaction between the two oppositely charged macromolecules and thus, enables us to measure the lifetime of the complex inside the nanopore. After showing that we can isolate a complex pair in the nanopore, we studied the effects of molecular identity on the nature of interaction in different complex pairs. In contrast to the irreversible conductance state of the alpha-hemolysin (αHL) channel in the complexation of poly-styrene-sulfonate (PSS) and poly-L-lysine (PLL), a reversible conductance state is observed during complexation between single stranded DNA (ssDNA) and PLL. This suggests that there is a weak interaction between ssDNA and PLL, when compared to the interaction in a PSS–PLL complex. Analysis of the PSS-PLL complexation events and its lifetime inside the nanopore supports a four step-mechanism: (i) The polyanion is captured by the pore, (ii) the polyanion starts threading through the pore. (iii) The polycation is captured, a complex pair is formed in the pore, and the polyanion slides along the polycation. (iv) The complex pair can be pulled through the pore into the trans compartment or it can dissociate. Additionally, we have developed a simple theoretical model, which describes the lifetime of the complex inside the pore. The observed reversible two-state conductance across αHL channel during ssDNA-PLL complexation, is described as the binding/unbinding of PLL during the translocation of ssDNA. This enables us to evaluate the apparent rate constants for association/dissociation and equilibrium dissociation constants for the interaction of PLL with ssDNA. This work throws light on the behavior of polyelectrolyte complexes in an electric field and enhances our understanding of the electrical aspects of inter-macromolecular interactions, which plays an extremely important role in the organization of macromolecules in the crowded and confined cellular environment.

---

Survival of living organisms relies on macromolecular assembly and their transport across cellular compartments in an electrolytic medium. For instance, protein import into mitochondria^1,2^, bacterial conjugation^3^, and transport through nuclear pore complexes^4, 5^. These processes occur repeatedly and energy is injected into the system by ATP, thus the biological system is driven away from equilibrium. Although nature’s biochemical machinery has inspired chemist to develop artificial molecular machines to mimic the form and function of their biological analogs^6^, we still lack the fundamental understanding of the non-equilibrium dynamics that occur in the confined and crowded cellular environment.

One of the simplest components of the biological system is a charged polymer in an electrolyte solution^7, 8^. There has been huge progress made in understanding the behavior of a single polyelectrolyte chain in an electrolyte solution^7, 8^. To take a step closer towards an understanding of the assembly of macromolecules and their dynamics in cellular environments, we must know how two oppositely charged polyelectrolytes interact with each other to form a complex pair. Apart from biological importance, polyelectrolyte complexes must be understood to design new materials^9, 10^ for separation science, flocculation, coating, and gene therapy applications^11,12^.

An experimental observation for the behavior of the polycation–polyanion complex in an electric field may give us valuable insight about the energetics of polyelectrolyte complexation. For example, the threshold electric field at which a complex dissociates may give us insight about the net interaction energy and the effects of the local dielectric constant on the interactions. However, if we simply mix two aqueous solutions of oppositely charged polymers, it is very difficult to isolate a polyelectrolyte complex pair as every macromolecule electrostatically interacts with many others. Additionally, to observe all the electrical aspects of the interactions, we must apply a sufficiently high electric field. In an electrolyte solution, an electric field can be induced by applying an electric potential gradient given by:

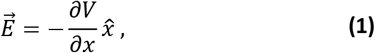

Where x^ is unit vector in the direction of the region of higher electric potential to the region of lower electric potential. Applying an electric potential gradient in a beaker across long distances will not create large enough electric field; To correct both issues we must isolate two oppositely charged polymers in close proximity to each other and we must apply the potential gradient across very narrow displacements (nm) such that induced electric field is high enough, and can generate stronger forces than the interaction forces in the complex pair. Fortunately, recent developments in nanopore technology have allowed for single-molecule analysis of macromolecules like DNA, RNA, proteins and synthetic polyelectrolytes^13–36^. Additionally, several studies have focused on manipulating the dynamics of polyelectrolytes passing through a nanopore^37–45^.

In this work, we propose using nanopore technology to drive a pair of oppositely charged macromolecules to form a complex. The confinement of nanopore in conjunction with an applied electric field provide the necessary precision to form an isolated polyelectrolyte complex. Our ionic current measurements across the pore identify a unique signature, which captures the dynamics of a polyelectrolyte complex under non-equilibrium conditions. Furthermore, we find that signatures of polycation-polyanion complex formation event can be distinguished for two different complex pair, PSS-PLL and ssDNA-PLL.

A schematic of our experimental setup is shown in Figure 1(a). We form phospholipid bilayer which creates two compartments, *cis*/*trans*, in our experimental chamber. To create a channel between chambers, we embedded a protein nanopore in the membrane. By loading polyanion in the *cis*, polycation in the *trans* chambers, and, by applying a voltage, we bias dynamics such that the oppositely charged polymers may meet inside the pore. This process can be monitored by measuring ionic current across the pore. When the macromolecule undergoes translocation through the pore, the ionic current is reduced to an average value of *I_b_* (occupied pore current) from the open pore current value of *I*_0_. Therefore, the ratio *I_b_*/*I*_0_ gives the state of pore while a molecule is passing through.

**Figure 1.**
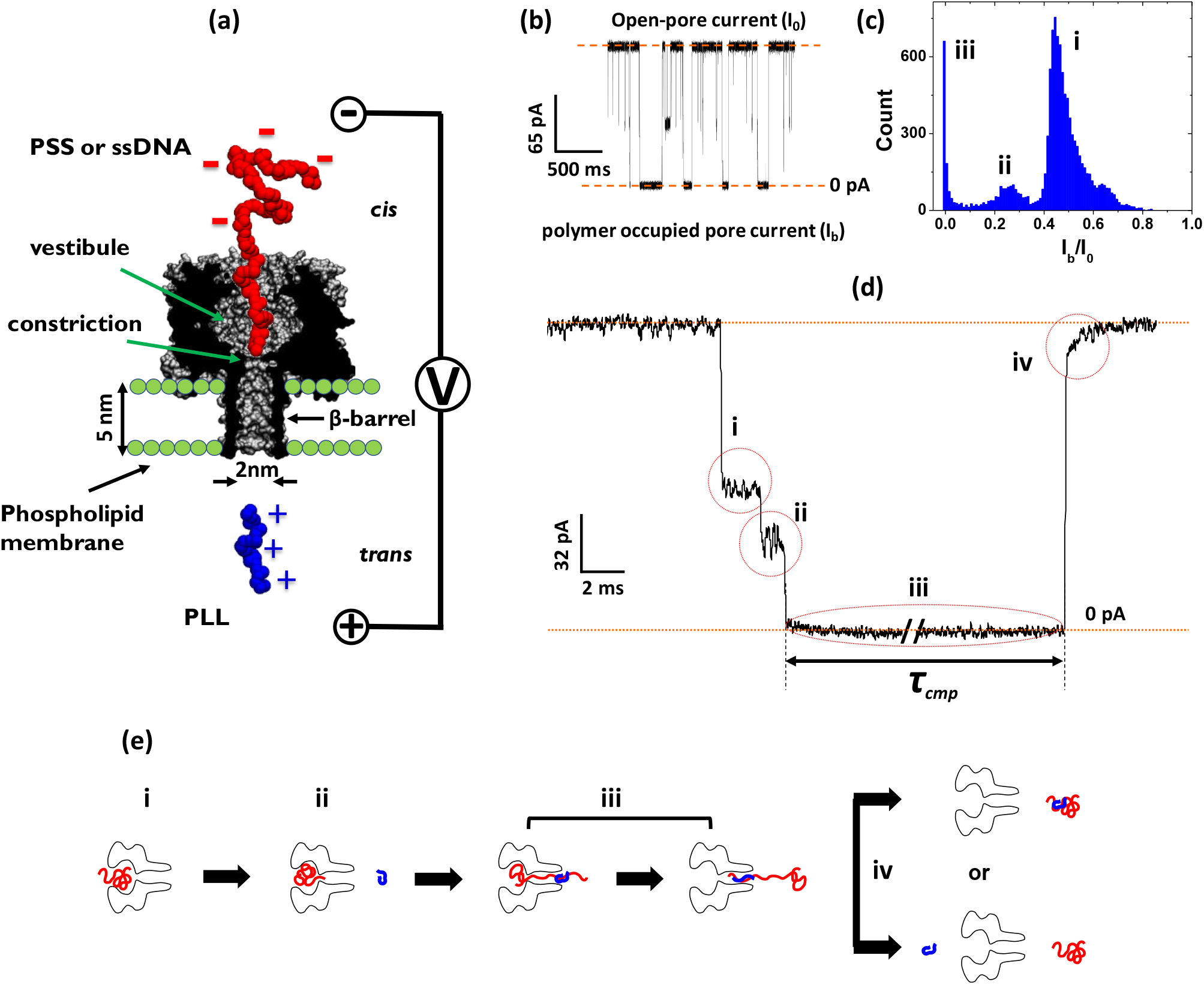
(a) Concept of voltage driven complexation in an α-HL pore, negatively charged NaPSS (red) on the cis side and positively charged PLL (blue) on trans side are subjected to electrophoretic capture by the same pore. (b) Representative ionic current traces when NaPSS and PLL were both subjected to electrophoretic capture. (c) Corresponding distribution of ionic current blockages (I_b_/I_0_). Three distinct peaks are observed in the distribution. The first two peaks centered on 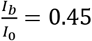 and 0.25 are identical with isolated NaPSS translocation events. The observed sharp peak at 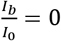 (pore is completely blocked), is attributed to complexation between NaPSS and PLL inside the pore. (d) Expanded view of ionic current levels in complexation between NaPSS and PLL. (e) Schematic of the observed four stages of complexation. All the measurements were carried out in 1M KCl, at7.5 pH and 30°C.

For the nanopore we employ the protein staphylococcal α-haemolysin (αHL) because of its well characterized nature. At its narrowest constriction, αHL is 1.5 nm in diameter, Sodium salt of polystyrene sulfonate (PSS) and single stranded DNA (ssDNA) were used as a polyanion and poly-L-lysine (PLL) as a polycation. PSS has been historically studied due to its electric and geometric resemblance to that of ssDNA with charge of −0.30e per monomer in 1M KCl solution and backbone thickness of 0.9 nm^8^. PLL has thickness of roughly 1 nm along its backbone, therefore only a single polymer can pass through the narrowest constriction (1.5nm) but not a complex pair. Previous work has established the ability of αHL pore, to distinguish between successful and unsuccessful translocations of PSS^13,35^. We take advantage of this capability to illuminate, for the first time, voltage directed complexation of two oppositely charged macromolecules. We performed all the experiments in 1M KCl, at 7.5 pH.

## Complexation of PSS and PLL

Before studying complex formation, we performed two control experiments; translocation of only PSS and translocation of only PLL. As described in earlier reports^13, 35^, for PSS there are three different types of events are observed during the translocation. At 140 mV, we observed three types of events with two main populations of I_b_/I_o_ (Supplementary info, SI: Figure S1). The duration of blockage which corresponds to I_b_/I_o_ < 0.35, is proportional to the molecular weight of PSS and inversely proportional to the voltage (SI: Figure S1 and S2). Therefore, only these blockages correspond to successful translocation events. For PLL, no clear translocation events are observed; only spike-like current blockages (duration less than 0.5 ms) were found corresponding to I_b_/I_o_> 0.40 (SI: Figure S3).

To observe complex pair formation, PSS was added to *cis* chamber, PLL was added to *trans* chamber, and a potential of 140mV to 240mV was applied. We observed different types of transient current blockages (Figure 1 (b) and (c)). In addition to the PSS events (i and ii), an event where the pore is nearly completely blocked (*I_b_*/*I*_0_ ≈ 0) is observed (iii). A representative current trace with such an event is shown in Figure 1(d), with very long blockage durations (in some cases longer than a minute) and four distinct current level indicating four stages (Figure 1(e)). The PSS enters the nanopore in vestibule (i) and gets conformationally trapped before reaching to the narrowest constriction (ii), upon entering narrowest constriction (β-barrel) trying to navigate through the pore, while PLL is approaching the pore from trans side (iii) PLL gets captured and makes complex with PSS which leads to complete blockage of the pore (i.e. *I_b_*/*I*_0_ ≈ 0) and depending upon strength of applied voltage PSS-PLL complex pair can be pulled through the pore or dissociate. In the following sections, we analyze each of these stages in detail.

## Stage i, polyanion capture in vestibule

The first stage is indicated by blockade of the open pore current (I_0_ = 162 ± 2.2 pA) to level i (Figure 1(d)), which has exactly same conductance value (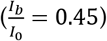) as isolated PSS in the vestibule of the pore (SI: Figure S (1)).

## Stage ii, polyanion enters into β-barrel

In next step the current drops further to level ii (Figure 1(d)), which has exactly the same conductance value (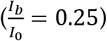), as PSS threading through narrowest constriction of the pore (SI: Figure S (1)), and into the β-barrel.

## Stage iii, complexation and sliding

The pore becomes completely blocked indicated by an ionic current that nearly vanishes (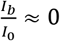, Figure 1(d)). We attribute this to complex formation between PSS and PLL molecules inside the nanopore. We rarely observe these types of events below 120 mV, which corresponds to the voltage threshold for successful PSS translocation. We also observe that populations of these completely blocked pore events depend upon molecular weight of PSS (SI: Figure S5). These observations suggest that complexation is occurs in the β-barrel. This hypothesis is further supported by the fact that the narrowest constriction of α-HL is too small to allow both PSS and PLL to pass through together.

To test this hypothesis more systematically, we denote these completely blocked pore events as complexation events and define a conditional probability as: 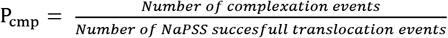; In this equation, we assume in every complexation event the PSS successfully translocates. We analyze the molecular weight and concentration dependences of this probability.

According to our hypothesis, P_cmp_ depends on both the PSS residence time inside the β-barrel and PLL capture rate. Therefore, the use of PSS with higher molecular weight and enhancement of PLL capture, will lead to a higher P_cmp_. Figure 2 shows the experimental measurements of P_cmp_. We found that indeed P_cmp_ increases with the NaPSS molecular weight (Figure 2 (a)) and is independent of NaPSS concentration (Figure 2(b)). Moreover, P_cmp_ is an increasing function of PLL concentration (Figure 2(d)) and shows weak dependence on PLL molecular weight (Figure 2(c)). Weak dependency of P_cmp_ on PLL molecular weight can be explained by the small changes in capture rates of PLL for the three different chain lengths employed. All the above results clearly validate that these long, completely clogged pore events are due to a complex pair formation between PSS and PLL in the β-barrel.

**Figure 2.**
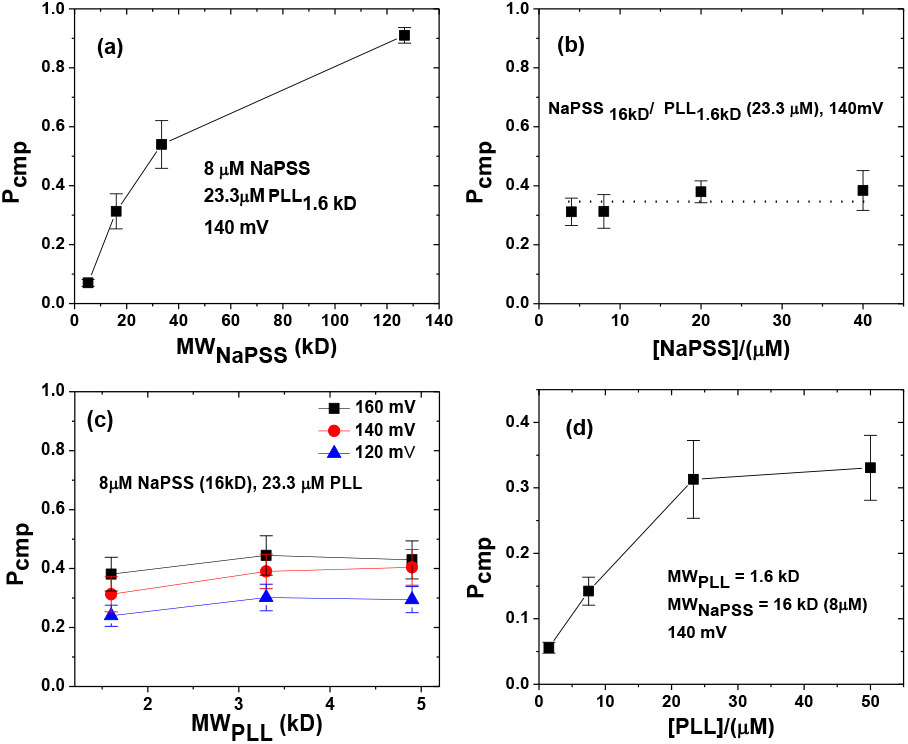
Probability of complexation (P_cmp_) event as a function of (a) Molecular weight of NaPSS (b) Concentration of NaPSS. (c) Molecular weight of PLL. (d) Concentration of PLL. Error bars represents standard deviation obtained from three independent measurements.

## Lifetime of complex inside the nanopore

To examine the dynamics of the complex pair, we measured the duration of complexation event (τ_cmp_) as shown in Figure 1(d). We describe, τ_cmp_ to be lifetime of the complex inside the pore, which characterize the complex pair dynamics. To interpret the complex-life time, we first fit the observed histogram of τ_cmp_ with the log-normal probability density function, and obtained the mean value of distribution represented by τ_cmp_. As shown in Figure 3(a), τ_cmp_ decreases exponentially as a function of applied voltage and increases with the molecular weight of NaPSS. These results clearly indicate that during complexation events (Fig. 1(d)), PSS must be pulled through the pore into the *trans* compartment. This further supports our assumption that in every complexation event PSS successfully translocate.

**Figure 3.**
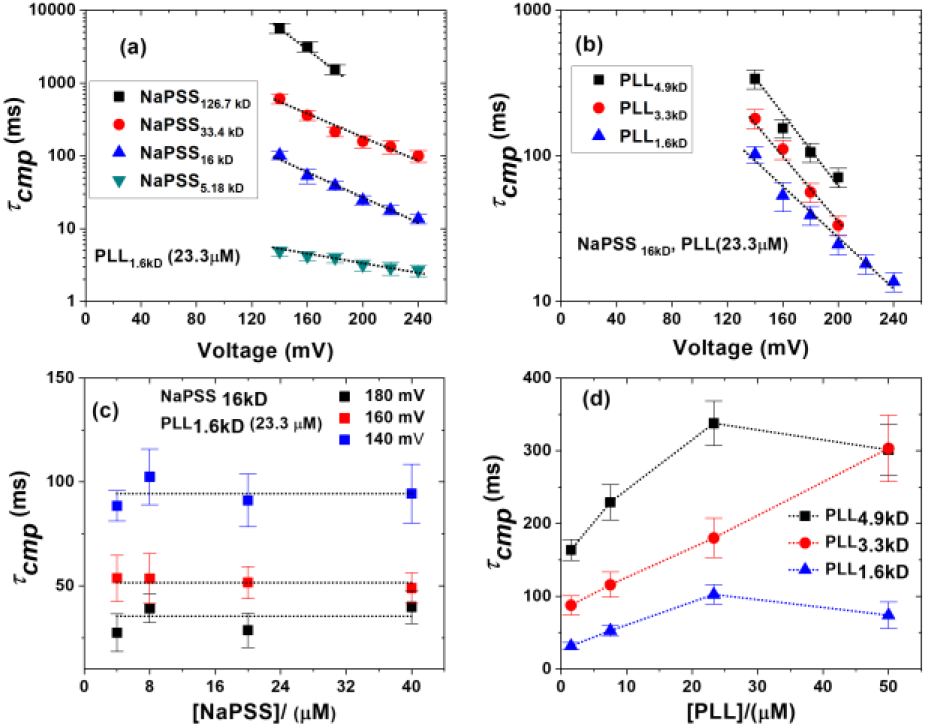
Most probable complexation event duration as a function of voltage for (a) different molecular weight of NaPSS, (b) different molecular weight of PLL. Dotted line represents exponential fit (c) Most probable complexation event duration as a function of concentration of NaPSS and (d) concentration of PLL. Error bars represents standard deviation obtained from three independent measurements.

As PSS is pulled into the *trans* chamber, it must slide along the PLL chain (Figure 1(e), iii). This sliding motion is accompanied by a friction due to electrostatic interactions between PSS and PLL. This sliding mechanism can be observed in an increase in τ_cmp_ with increasing molecular weight of PLL (Figure 3(b)). As longer PLL molecule can create more electrostatic frictional force against PSS translocation. The value of τ_cmp_ does not depends on the concentration of PSS, but increases with concentration of PLL (Figure 3(c) and 3(d)). The former fact aligns with intuition that complexation occurs with a single molecule of PSS and PLL in the pore. The increase in τ_cmp_ with PLL concentration can be understood because capture of PLL occurs more quickly leaving the complex more time to remain in the pore. However, we also observe that the life–time of the complex in the pore behave differently for different molecular weight of PLL (Fig.3 (d)). The exit ring of the β-barrel has 7 negative charges, which can also interact electrostatically with PLL, and can make PLL to be fixed in the pore while PSS is sliding.

## Stage iv, complex pair translocation or dissociation

Based on information that in every complexation event, PSS is successfully translocated via sliding along PLL, there can be two possibilities (Fig. 1(e)). One possible case is that the whole PSS-PLL complex pair is pulled out of the pore into the *trans* compartment. This is likely the case when the applied voltage is not large enough to break the complex pair apart. When the applied voltage is above a threshold, PLL will dissociate with PSS and move into *cis* compartment (as illustrated in Fig. 1(e)). Given, in 90% case of the complexation events, the blocked pore current directly recovers to the open pore current (iii→iv Fig. 1(d)), it is more likely that PSS-PLL complex pair translocate to *trans* side. If any complex pair dissociation is occurring in the pore, then one would expect to see at least one current level in between iii and iv (Fig. 1(d)). However, it could also be possible that after the dissociation of the complex pair, the dwell time of the individual polyelectrolytes in the pore are so small that we cannot measure it in given experimental resolution.

To describe the strong dependence of life-time of the complex in the pore on voltage, molecular weight of PSS and PLL, we developed a simple analytical model for our system. We follow the theory of reference (46,47 and 48), which describe the free energy of negatively charged polymer translocating through a nanopore. The theory predicts the free energy and translocation time with respect to two parameters:(i) effective charge on individual monomer of the polymer (q) and (ii) effective pore-polymer interaction (ε). The presence of PLL inside the β-barrel can be modeled (Fig. 4 (a)) as a pore having two different pore-polymer interactions (ε_1_ and ε_2_). This can be further manifested by presence of negatively charged amino acid residues at the end of β-barrel, which can help PLL to bind with the pore. Therefore, creating a net pore having two distinct regions with different pore-polymer interactions. Although, based on ionic current traces observed in experiment, there can be possibility of complex pair translocation, this model assumes that complex pair dissociates. We also assume that the electric potential drops linearly across the pore ^46,47,48^.

**Figure 4.**
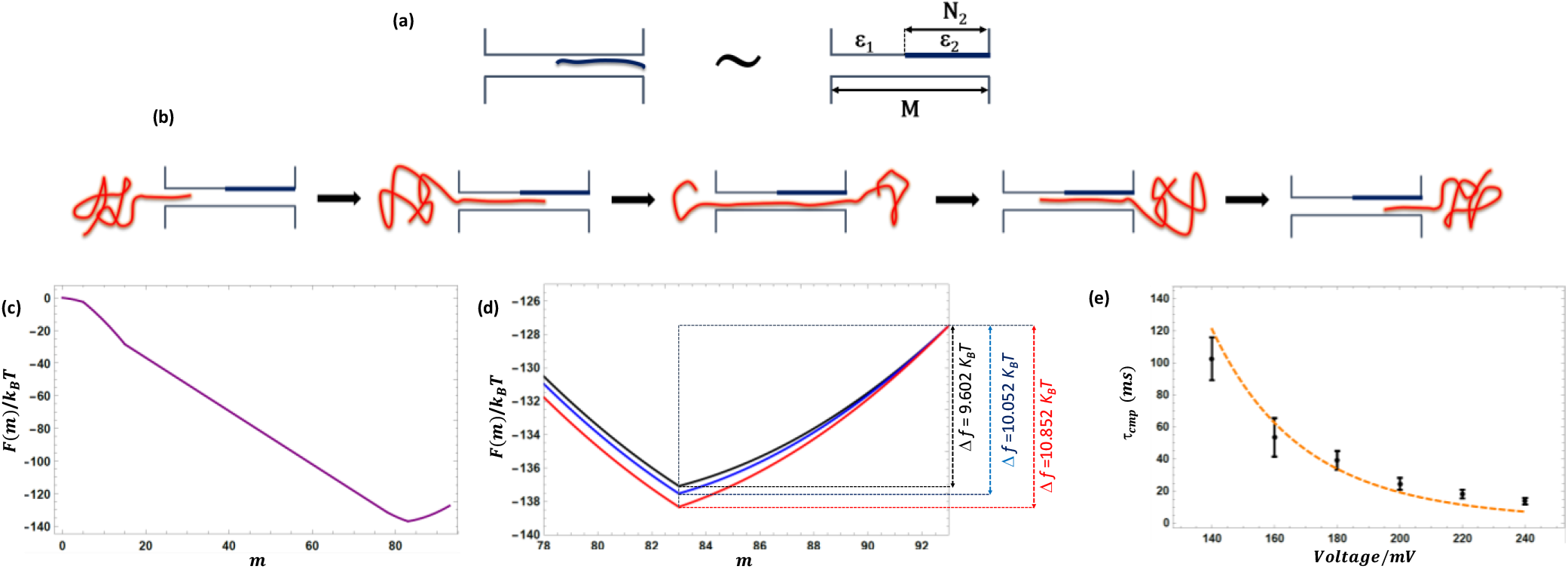
**(a)** Theoretical model describing the presence of positively charged PLL inside the β-barrel as a pore having two different regions, interacting differently with translocating negatively charged polymer (NaPSS). **(b)** Five stages in the translocation of NaPSS through the pore. **(c)** A representative free energy landscape for the translocation process as a function of translocation co-ordinate m. ε_1_ = 0.25, ε_2_ = 1.5, q=0.30, N_1_=78 (33.4kD NaPSS) and N_2_=10 (1.6 kD PLL). **(d)** Free energy landscape during the ejection stage of the polymer (l6kD NaPSS) for PLL_1.6kD_ (Black, ε_2_ = 1.505), PLL_3.3kD_ (Blue, ε_2_ = 1.550) and PLL_4.9gkD_ (Red, ε_2_=1.630). ε_1_ = 0.05, q=0.30, N_1_=78, N_2_=10 **(e)** Experimentally observed and calculated values of τ_cmp_ are plotted together. The black dot with error bars represents experimental data for the 16 kD NaPSS and 1.6kD PLL. The orange dash curve represents fit from the model (ε_1_= 0.05, ε_2_=1.505, q=0.30, N_1_=78, N_2_=10).

The translocation of PSS through a nanopore having regions of two different pore-polymer interaction consists of five stages (shown in Fig. 4(b)). We derive the free energy landscape for the translocation process. For an effective charge q on individual translocating monomer, N_1_ monomers of PSS, N_2_ monomers of PLL and M pore length, the free energy landscape for the translocation process with respect to translocation coordinate m is given equation 2

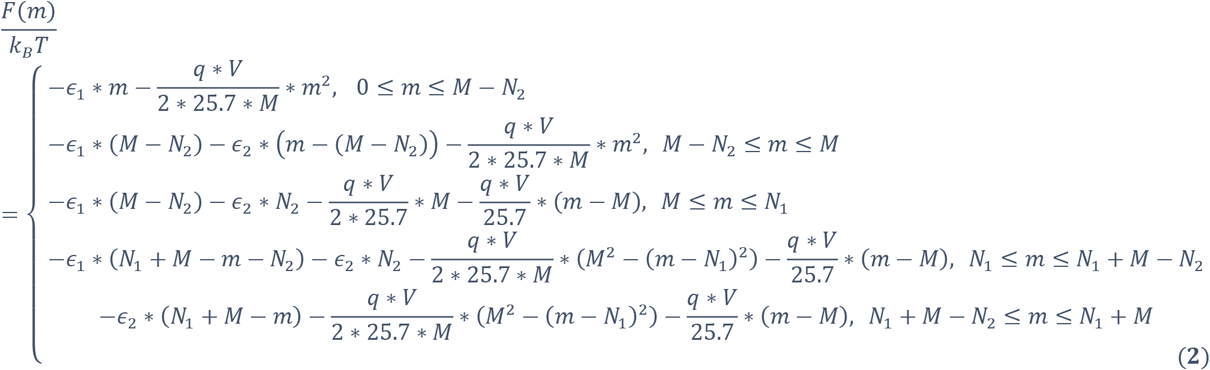

where *k_B_* is boltzman constant and T is temperature and *k_B_T*/*e* is taken as 25.7 mV at room temperature and e is charge of an electron. For such strong electric fields (20-50 mV/nm) in our experiments, the contribution from conformational entropy of the polymer outside the pore is negligible and thus is ignored in the above equation. For q=0.30 (Based on the manning argument), ε_1_ = 0.25 and ε_2_=1.5 a typical landscape is given in Fig. 4(c), where *V* =140*mV*, *M* = 15(corresponding to the length of β-barrel 5 nm), *N*_2_= 10 (PLL 1.6kD) and *N*_1_= 78 (corresponding to 16kD NaPSS). The role of the attractive electrostatic interaction between the PSS and PLL is manifested as an attraction between PSS and pore, which lead to a barrier (about 15*k_B_T*) in the last stage of the translocation.

We calculated the *τ_cmp_* using free energy of last three stages (m>M). We employ the theoretical machinery most suited to connecting the free-energy landscape and the time evolution of the probability of realizing a state *W_m_* (*t*) is the Fokker-Planck formalism. In general, the Fokker-Planck equation for the situation presented here is given by equation 3 as,

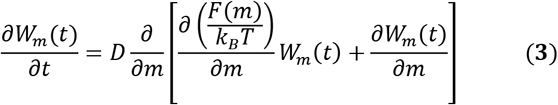

where D is the monomer diffusion coefficient related to local friction inside the pore ^(46,47,48)^. Substitution of Eq. 1 into Eq. 2 gives the time dependence of the probability of realizing a value of m in the reaction coordinate. Using standard procedures in the Fokker-Planck formalism, the average time, *τ_cmp_*, for crossing state of M to *N*_1_+*M* is given by equation

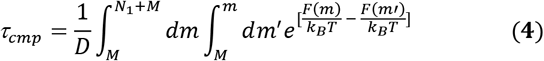

where *F*(*m*) is given by Eq. 1. In defining *τ_cmp_*, we used the reflecting boundary condition at m=M and the absorbing boundary condition at m=*N*_1_, + *M*, which are evident from our experimental observation as, the polyanion very rarely retracts back into the donor compartment after some monomers are threaded through the pore.

This model describes the voltage response of the life-time of the complexation events for all molecular weight of NaPSS and PLL employed with ε_2_ as an adjustable parameter (Fig. 4(d), 4(e) and SI: S7-S14). Increasing the molecular weight of NaPSS and PLL can be modeled as an increase in the net electrostatic attraction with the pore (ε_2_). An increase in molecular weight of NaPSS and PLL may lead to an increase in ε_2_ and therefore increase in the energy barrier in the landscape during the ejection of negatively charged polymer. A list of ε_2_ and energy barrier during the ejection are given in Table 1 obtained from the best-fit for the *τ_cmp_*, calculated from the model and observed in the experiments.

**Table 1.** Values of interaction energy parameters ε_2_ and free energy barrier Δf obtained as a result of best fit of calculated τ_cmp_ from the model and experimentally observed τ_cmp_.

| $N_{PLL}$ | $N_{PSS}$ | $\epsilon_2(k_B T)$ | $\Delta f(k_B T)$ |
| --- | --- | --- | --- |
| 10 | 25 | 1.115 | 5.702 |
| 10 | 78 | 1.505 | 9.602 |
| 10 | 162 | 1.731 | 11.862 |
| 10 | 615 | 1.965 | 14.202 |
| 20 | 78 | 1.550 | 10.052 |
| 30 | 78 | 1.630 | 10.852 |

**Table 2.** Table for the kinetic and apparent equilibrium constant for different length of poly-L-lysine measured at 120 mV.

| $k_{on} (\text{s}^{-1}\mu\text{M}^{-1})$ | $k_{off} (\text{s}^{-1})$ | $K_d^{app} (\text{mM})$ | PLL Length |
| --- | --- | --- | --- |
| $0.49 \pm 0.08$ | $1937.85 \pm 148.61$ | $3.93 \pm 0.71$ | 10 |
| $0.28 \pm 0.07$ | $419.02 \pm 138.76$ | $1.50 \pm 0.61$ | 20 |
| $0.37 \pm 0.07$ | $297.03 \pm 21.08$ | $0.79 \pm 0.15$ | 30 |

## Complex lifetime based on double exponential probability density function

Although the log-normal fits of the τ_cmp_ histograms provide the valuable insight and qualitative behavior of the complex lifetime, it is insufficient for understanding the underlying physical processes and thus making quantitative predictions. A physical basis for the choice of log-normal probability density function for fitting is also absent. Therefore, in this section, to enhance our understanding about observed histogram of τ_cmp_, we describe complex lifetime based on a double exponential probability density function. A physical basis for the choice of this function is provided in supplementary information (section (vi)).

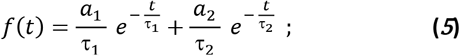

We find good agreement of complex life-time histogram with double exponential forms (SI: S15, S16, and S17); indicating two underlying physical processes, occurring with time constant τ_1_ and other with time constant τ2. To understand the both time constants, we analyzed their dependency with respect to molecular of PSS and PLL respectively at different voltages.

We find that both time constants increase with molecular weight of PSS ((Figure 5(a), and 5 (b))) and PLL (Figure 5(c), and 5 (d)), which is in qualitatively agreement with the fit of complex life-time based on log-normal probability density function. Interestingly, the slower time constant τ_2_ seem to behave quadratically with molecular weight of PSS. However, τ_1_ seem to slightly deviate from quadratic behavior. These observations, support the common sliding mechanism in dynamics of complex pair and opening a new door for further theoretical and computational investigations. Another interesting observation is that for almost every condition the slower time constant τ_2_ is 10 times than that of faster time constant τ_1_, i.e. 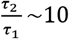. We also find that the behavior of τ_1_ and τ_2_ with respect to voltage can be justified based on the theoretical model described in previous section (SI: S18, and S19). The parameters obtained from the fit with the theoretical model is given in supplementary information ((SI: S20, and S21)). In most of condition the slower time constant seem to have more contribution compare to the faster time constant (SI: S22).

**Figure 5.**
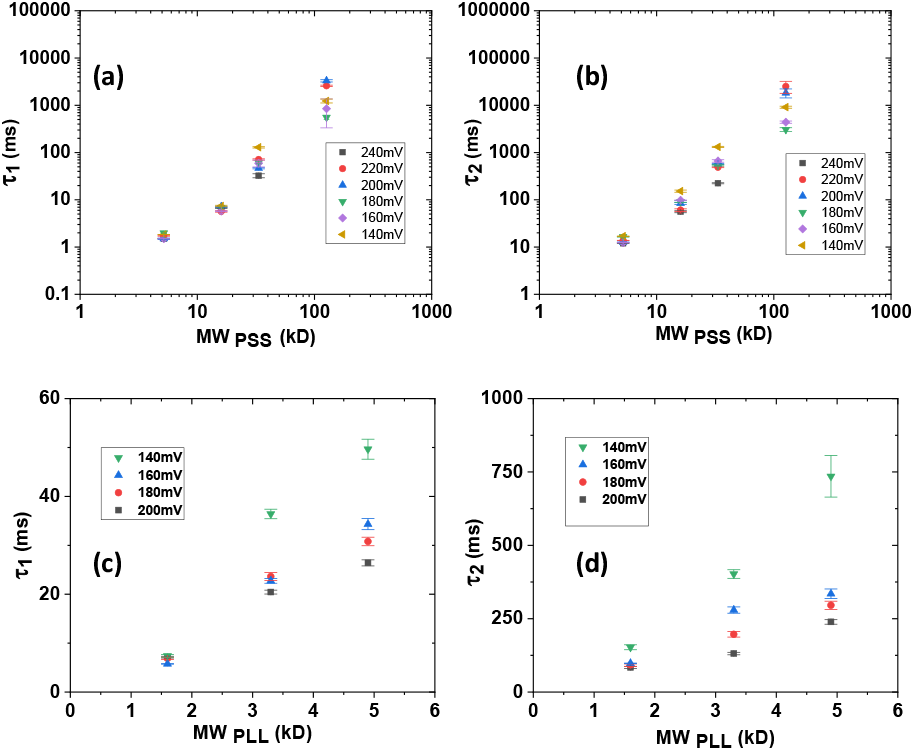
Effect of molecular weight of NaPSS on time constant **(a)** τ_1_ for 23.3 μM 1.6kD PLL **(b)** τ_2_ 23.3 μM PLL1.6kD. Effect of molecular weight of PLL on **(c)** τ_1_ and **(d)** τ_2_ at different voltages (23.3 μM. Error bars represents standard deviation obtained in fitting the histograms with double exponential probability density function.

## Complexation of ssDNA and PLL

In this section, we use this new approach of voltage-driven polyelectrolyte complexation to determine whether and how intermolecular interactions in a complex pair of negatively charged ssDNA with positively charged poly-L-lysine (PLL), differs from that of interactions in complex pair of negatively charged polystyrene sulfonate (PSS) with positively charged PLL. Although, PSS has been celebrated as a model system to understand the behavior of negatively charged bio-polymer in the electrolytic solution due to its geometric and electrical resemblance to that of ssDNA^1–6^, the location of the charge on the backbone and local dielectric constant may differ and therefore interactions of these two negatively charged polymer (ssDNA and PLL) with positively charged polymer may be different.

In contrast to complexation of PSS-PLL in the pore, for ssDNA-PLL complexation we observed a unique type of events where ionic current undergoes a reversible transition between the two states (Figure 6 (a) and (b)). Suggesting, the effect of local dielectric heterogeneity due to the specificity of the macromolecules and there is weaker interaction in ssDNA-PLL complex pair compared to the PSS-PLL complex.

**Figure 6.**
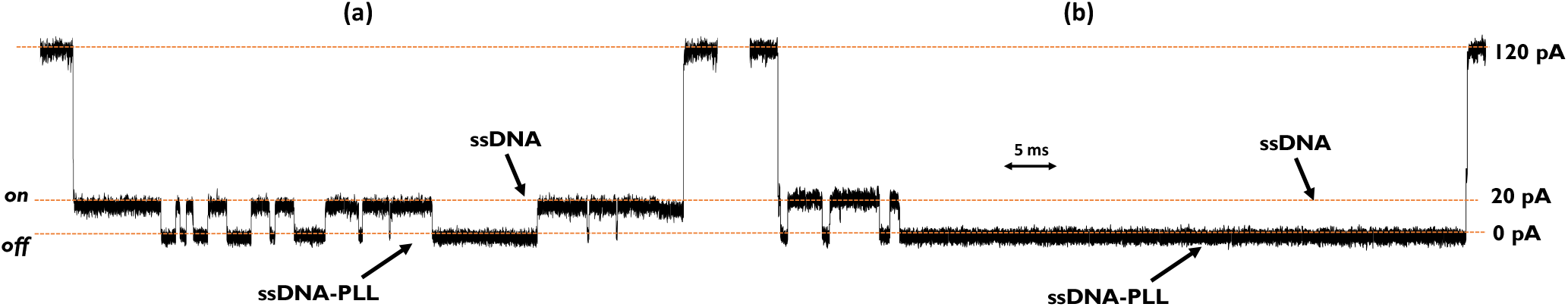
Some representative ionic current traces for the ssDNA-PLL complexation events for PLL3.3kD at 120mV.

Before studying the interaction between ssDNA and PLL using *α*HL pore, we performed two control experiments: translocation of only ssDNA and translocation of only PLL. At an applied voltage of 120 mV during ssDNA (*cis*) translocation, open pore current (120pA) blocked to less than 20pA (SI: S23 (a)) and the most probable translocation time for ssDNA is 0.17 ms, which agrees with the earlier reports^7^ for the same sequence of ssDNA (SI: S24). We also observed some of the events where the current blocked for a longer time and blocked current is near about 20pA (SI: S23 (a)). For PLL induced blockage, we observed most of the events correspond to nearly 50 % blockage of open pore current, with (SI: S23 (b)). However, most of the durations were short (less than 1 ms) (SI: S25). We also observed some events which last for a longer time (For PLL3.3kD and PLL4.9kD) where current is blocked to 35±5 pA and which might be due to the binding of PLL with the β-barrel of the pore (not shown).

To detect the interaction between ssDNA and PLL, ssDNA was added to *cis* chamber, PLL was added to trans chamber, and an electrical potential of 120mV to 180mV was applied such that the two molecules may meet inside the pore, during this process we observed different types of transient current blockages. In search for the signature of interaction between ssDNA and PLL, we observed a type of event where the open pore current blocked to the level where the ionic current is identical to ssDNA translocation (20 pA), and then further blocked reversibly to the level where ionic current nearly vanishes as shown in Figure 6(a) and 6(b).

## Comparison of conductance state of the pore in ssDNA-PLL and PSS-PLL complexes

In the complexation of PSS-PLL, we learnt that when PLL molecule meet translocating PSS inside the pore, the ionic current nearly vanishes, however, current does not go back to the level of PSS translocation. In other words, in PSS-PLL complexation events, conductance state of the pore is irreversible, where as in ssDNA-PLL, the ionic current undergoes transition between two reversible states. The reversible transition between two conductance states may be due to the binding/unbinding of PLL with translocating ssDNA. The reversible nature of binding of PLL with ssDNA is may be due to the location of charges on ssDNA, which are protected (Figure 7), causing weaker electrostatic interaction. In contrast, charges in PSS are relatively exposed to the PLL therefore interactions can be stronger and therefore we have irreversible conductance state which involves sliding of PSS over PLL (Figure 1(e)).

**Figure 7.**
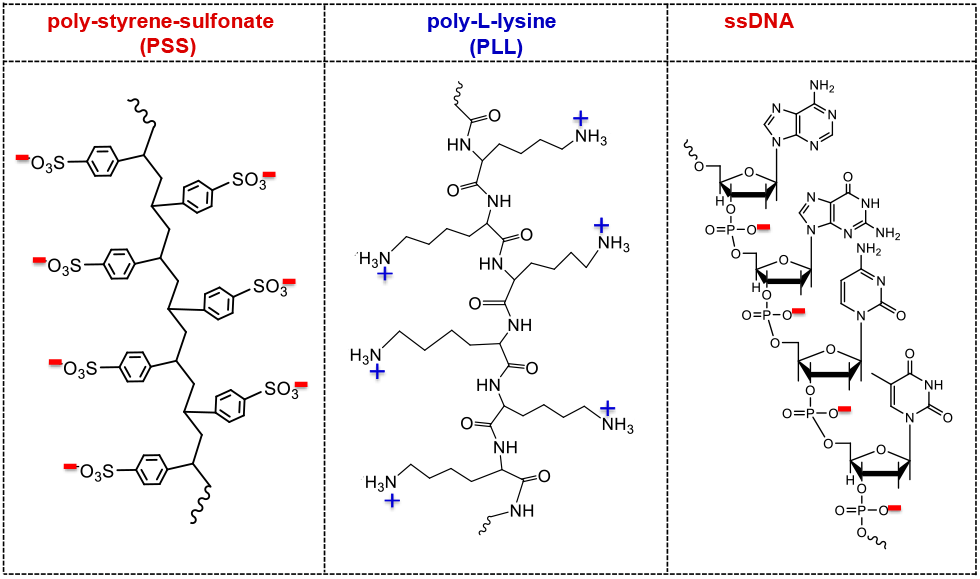
A sketch of chemical structures of PSS, PLL, and ssDNA.

We also observed that there is a distribution of number of events undergoing reversible transition between the two states (SI: S26), with mean value obtained from fitting the histogram with log-normal function, decreases by increasing PLL molecular weight.

## A two-state (*on*/*off*) model for ssDNA-PLL interaction

We hypothesized that the first ssDNA gets captured and while trying to navigate through the pore, PLL gets captured and bind with ssDNA, which leads to a complete blockage of the pore, PLL dissociate leaving only ssDNA in the pore and then again binds with ssDNA. This reversible process continues to occur until ssDNA is pulled out of the pore completely. We analyze these reversible transitions between the two states as a binding/unbinding of PLL with translocating ssDNA. We describe the state of the pore when ssDNA is passing through the pore as “*on*” state and when ssDNA bind PLL as “*off*’ state, we define a two-state model corresponding to the binding/unbinding of PLL with ssDNA (Figure 8). To validate our hypothesis, we first constructed the histograms for *on* state lifetime (τ_on_) and *off* state lifetime (τ_off_) at various voltages and concentrations of PLL.

**Figure 8.**
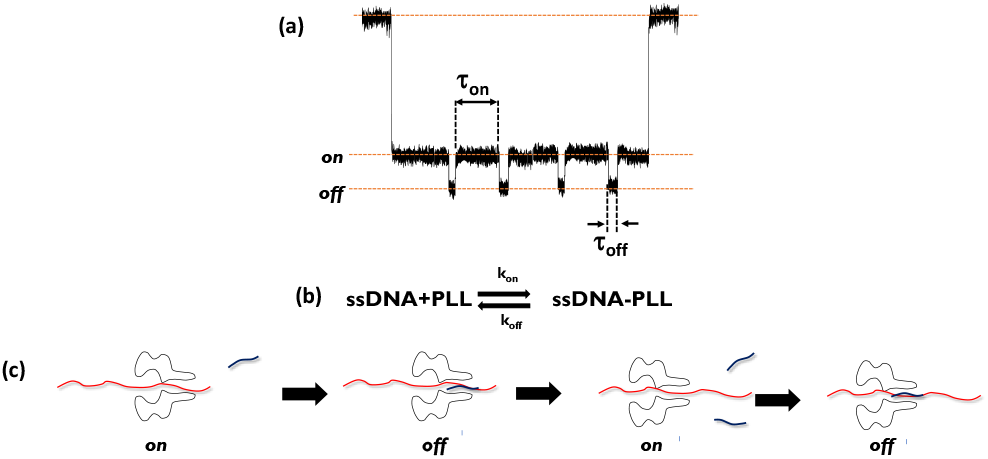
**(a)** A representation of on and off conductance sates corresponding to binding/unbinding of PLL with translocating ssDNA. **(b)** Binding/unbinding equilibrium scheme for ssDNA-PLL interaction. **(c)** Hypothesis of observed reversible conductance state as a binding/unbinding of PLL molecule with translocating ssDNA.

Histograms displaying the dwell time for the *on* and *off* state could be fitted by single-exponential, giving a time constant, which is nearly same as time constant obtained from the fit of histograms with log-normal probability density function (Figure 9 (a), 9(b), 9 (d), and 9(e)). These results agree with the existing binding/unbinding models for the bimolecular interactions^51–54^. Single time constants for τ_on_ and τ_off_ affirm our two state-model for the process and suggest that, under the conditions examined here, there exist only one mode for the interaction of ssDNA and PLL. Therefore, the kinetics of the interaction between ssDNA and PLL should obey the simple kinetic scheme given in the model (Figure 8(b)).

**Figure 9.**
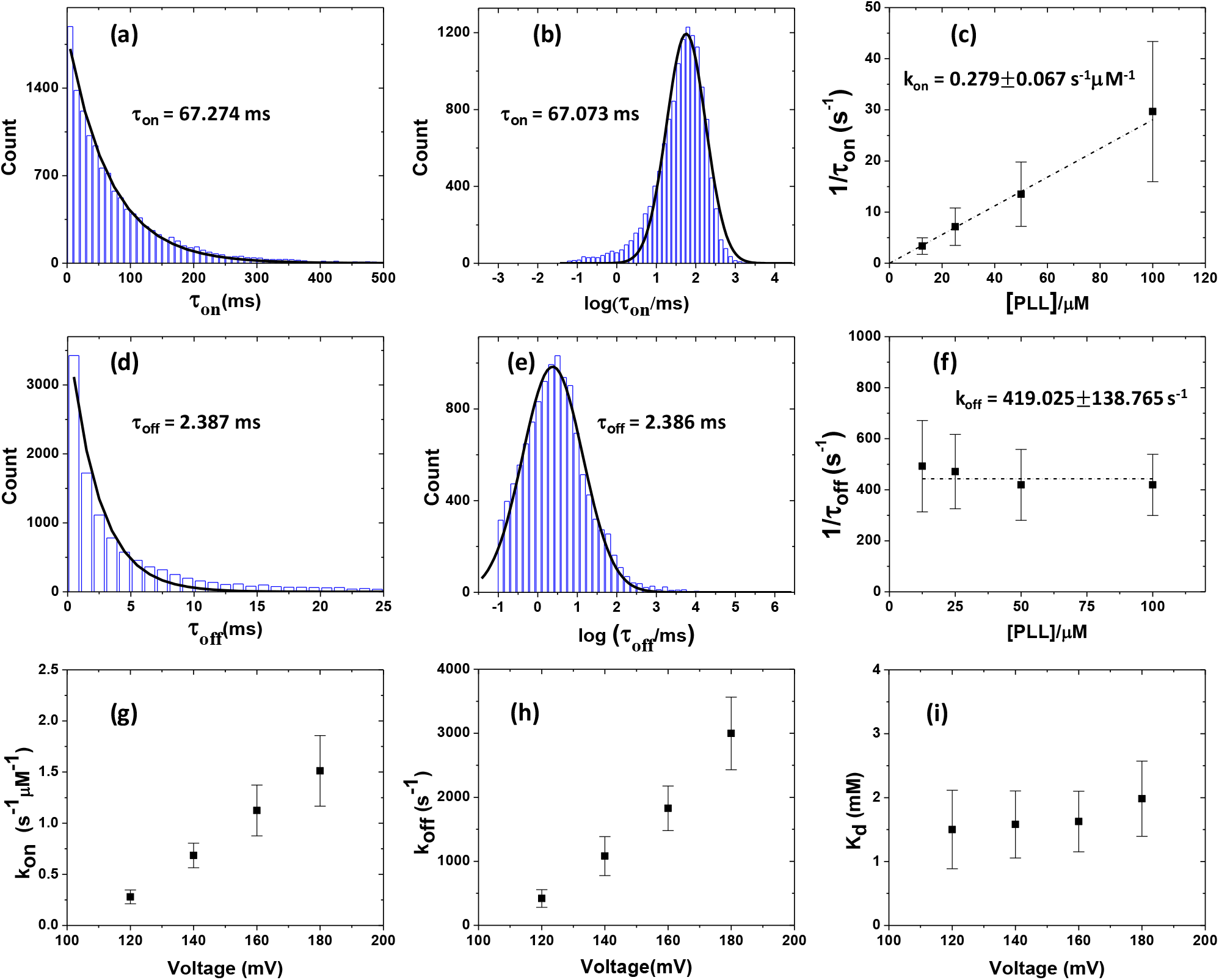
**(a)** Histogram displaying dwell times of “on” state (association time) at 120mV,for PLL_20_. The black curve represents single-exponential fit to the histogram, revealing a time constant of 67.274 ms **(b)** Histogram of logarithmic of dwell times if “on” state. The black curve is a log-normal distribution fit to the histogram, giving a time constant of 67.073 ms. **(c)** The rate constant k_on_ for the association between ssDNA and PLL is obtained from the slope of the linear fit to 1/τ_on_ vs [PLL]. **(d)** Histogram displaying dwell times of “off’ state (dissociation time) at 120mV, for PLL_20_. The black curve represents single-exponential fit to the histogram, revealing a time constant of 2.387 ms **(e)** Histogram of logarithmic of dwell times if “off’ state. The black curve is a log-normal distribution fit to the histogram, giving a time constant of 2.386 ms. **(f)** The rate constant koff for the dissociation of ssDNA-PLL complex is obtained from the average of 1/τoff values. **(g)** The dependence of association rate constant k_on_ with voltage, suggesting electric field kinetically favor the association **(h)** The dependence of dissociation rate constant k_off_ with voltage, suggesting electric field kinetically favor the dissociation **(i)** The dependence of equilibrium dissociation constant is nearly independent with voltage, suggesting electric field is thermodynamically inert to binding/unbinding of PLL with ssDNA. Error bars represents standard deviation obtained from three independent measurements.

As expected from our two-state model, the plot of 1/ τ_on_ is a straight line with respect to concentration of PLL, the slope of which yields the rate constant *k*_on_ for association between ssDNA and PLL (Figure 9 (c)). Again, in agreement with our two-state model, 1/ τ_off_ is independent of [PLL] (Figure 9 (f)) and yields a rate constant *k_off_* for the dissociation of ssDNA-PLL, which is evaluated as the average of 1/ τ_off_ at the different PLL concentrations.

## Voltage-dependency of *on*/*off* kinetics

By measuring τ_on_ and τ_off_ at various concentration of PLL for different voltages, we analyzed the rate constants for the association and dissociation of ssDNA-PLL complex. The association rate constant *k*_on_ increases with applied transmembrane voltage (Figure 9(g)), suggesting that voltage, kinetically favors the association of ssDNA and PLL. The dissociation rate constant *k*_off_ also increases with applied transmembrane voltage (Figure 9(h)), indicating that dissociation of ssDNA-PLL is also kinetically favored by the voltage. We have also analyzed the apparent equilibrium dissociation constant (K_d_^app^= *k*_off_/*k*_on_) defined as the ratio of *k*_off_ and *k*_on_. Surprisingly, the value of K_d_^app^ does not depend upon voltage (Figure 9(i)), and value around 1.6 mM indicates that the interaction between ssDNA and PLL is weak. This observation indicates that the voltage is thermodynamically inert to the association and dissociation between ssDNA and PLL molecules.

## Effect of PLL length on *on*/*off* kinetics

We have also investigated the effect of PLL length on the kinetics of association/dissociation of ssDNA-PLL. The association rate constant *k*_on_ depends weakly upon length of PLL. At voltage of 120 mV, for PLL_10_ (Mw = 1.6kD) the value of *k*_on_ is 0.49 ± 0.08 s^−1^μM^−1^. The values are 0.28 ± 0.07 s^−1^μM^−1^ and 0.37 ± 0.07 s^−1^μM^−1^ for PLL_20_ (Mw = 3.3kD) and PLL_30_ (Mw = 4.9kD) respectively. Higher PLL length diffuses faster, therefore one will expect to have higher association rate constant for the shorter PLL, if association is diffusion limited. On the other hand, longer PLL contains more number of charges, and if one assumes that the interaction between ssDNA and PLL is purely electrostatic, a longer PLL may have higher attraction with ssDNA. Therefore, longer PLL may have a higher association rate constant if association is attraction limited. The observed *k*_on_ with respect to PLL length may have both effect leading to non-monotonicity in *k*_on_.

The dissociation rate constant *k*_off_ decreases with increasing length of the PLL. At 120 mV, for PLL_10_ the value of *k_off_* is 1937.85 ± 148.61 s^−1^. The values are 419.02 ± 138.76 s^−1^ and 297.03 ± 21.08 s^−1^ for PLL_20_ and PLL_30_ respectively. If the dissociation is diffusion limited then one would expect to have faster dissociation rate constant for shorter PLL. If the dissociation is against the electrostatic interaction between ssDNA, then also one would expect to have higher dissociation rate constant for shorter PLL. As longer PLL may get arrested by ssDNA due to more electrostatic attraction. Therefore, observed *k*_off_ align with our intuition. The non-monotonic dependence of *k*_on_ and monotonic dependence of *k*_off_ indicate us that electrostatic interactions playing a major role in this process.

The equilibrium dissociation constant K_d_^app^ decreases with increasing the PLL length. At 120 mV for PLL_10_ the observed value of K_d_^app^ is 3.93 ± 0.71 mM, the values are 1.50 ± 0.61 mM and 0.79 ± 0.15 mM for PLL_20_ and PLL_30_ respectively. These observations manifesting the role of number of charges present in the system and align with the intuition that longer PLL will be harder to dissociate because of strong electrostatic interaction. These observations indicate that the PLL acts as a multivalent cation in the system.

## Summary

We have demonstrated that how oppositely charged macromolecules can be driven by electric field to form a polyelectrolyte complex inside a nanopore. Our ionic current measurements across the pore identify a unique signature of complex formation, which captures the dynamics of a complex pair in non-equilibrium condition. In total, we observe four stages in the complexation between PSS and PLL, and we have interpreted a macromolecular description for each. Our analysis of complexation events suggests that, after complexation in the pore, the PSS slides along PLL against a frictional force due to electrostatic attraction. Sliding phenomenon of SSB proteins over ssDNA has been recently studied^49, 50^, where protein sliding over DNA is described by reptation mechanism. Detection of signature and a theoretical description for complex dissociation are future scope of this work. Our simple analytical model describes the life-time of the complex inside the pore and indicates that small changes in interaction between PSS and PLL can account for substantial increase in life-time. The distributions of the measured complex lifetimes under various experimental conditions follow double–exponential form and thus it can be concluded that two distinct processes occur while the complex is inside the pore. The time scales of the two processes were obtained by fitting our collected data, and we found that one of the processes occurs an order of magnitude faster than the other.

The ionic current signature obtained in the complexation of ssDNA and PLL molecules, suggest there is a weak interaction between ssDNA and PLL when compared to the interaction in a PSS–PLL complex. In contrast to the irreversible conductance state in the PSS–PLL complexation measurement, during ssDNA–PLL complexation we observed a reversible conductance state, which enabled us to measure the rate constants for association and dissociation of PLL with translocating ssDNA and hence apparent equilibrium binding constants. The weaker interactions in ssDNA–PLL may be due to nature of the local dielectric constant; the negative charges on PSS are on a side group and are thus relatively exposed, whereas in ssDNA the charges are on the backbone and are more hidden. Due to this, the local dielectric constant near the charges is lower in PSS compared to ssDNA and so the electrostatic interactions in the PSS–PLL complex are stronger than in the ssDNA–PLL complex. This conclusion can be further manifested by comparing the translocation time of ssDNA and PSS. Under identical conditions, the same number of monomer of PSS translocate through the αHL pore in 1 ms, whereas ssDNA takes only 0.2 ms for translocation. This is because the charge is exposed in PSS, causing more counter ion condensation and reducing the electrophoretic force on it. On the other hand, in ssDNA charge is relatively hidden, causing less counterion condensation and a higher electrophoretic force, therefore it moves faster through the pore.

Measurements of lifetime of complex at different temperature, for molecular identity, different nanopores, and at higher voltages are future scope of this work.

## SUPPORTING INFORMATION

Materials, methods, additional data for control experiments and complexation experiments. This material is available free of charge via the Internet at http://pubs.acs.org.

## ACKNOWLEDGMENT

Acknowledgement is made to the National Science Foundation (Grant No. DMR 1404940) and National Institutes of Health (Grant No. R01HG002776-11).

