## Supplementary Information for "Voltage-driven polyelectrolyte complexation inside a nanopore"

This supplementary information contains the following:

(i) Materials/Methods.

(ii) Control experiments for NaPSS only translocation and PLL only translocation (Figure S1-S4).

(iii) Additional data for PSS-PLL complexation (Figure S5-S6).

(iv) Free energy landscape during ejection of the polymer as described by model in main text for different molecular weight of NaPSS (Figure S7-S9).

(v) Fit of the experimental data with the model described in text (Figure S10-S14).

(vi) Analysis of complex life-time based on double exponential probability density function (S15-S22).

(vii) Experiments for ssDNA-PLL complexation (Figure S23-S31).

### (i) Material and Methods

**Formation of perpendicular bilayer:** We used 1,2-diphytanoyl-*sn*-glycero-3-phosphocholine (DphPC, purchased from Avanti Polar Lipid.) to form perpendicular phospholipid bilayer across an aperture (diameter, 50 μm) in a Teflon film separating the two compartments (1 ml each *cis*/*trans*) of the polytetrafluoroethylene (PTFE) flow cell apparatus (purchased from Nanopore Solutions, Portugal).

**Buffer solution:** 1M potassium chloride (KCl) solutions in 10 mM 2-[4-(2-hydroxyethyl) piperazin-1-yl] ethanesulfonic acid (HEPES) in deionized water (18 MΩcm) were prepared for measurements. The pH of the buffer solution was adjusted to 7.5 by adding potassium hydroxide (KOH) or hydrochloric acid (HCl).

**αHL pore:** Wild-type αHL monomers, α-toxin from staphylococcus aureus were purchased from EMD Millipore (MA). A stock solution of αHL were prepared by adding 50:50 (v/v) of water and glycerol mixture to make final concentration of 0.5 mg/ml. This resulting solution was further diluted to by 1M KCl buffer solution to make final concentration of 0.5 μg/ml. A portion of the resulting αHL protein solution (2 μl) was added to the *cis* compartment of a bilayer apparatus to form a single-channel. Once the single pore is formed, the *cis* chambers is flushed with 1M KCl solution to avoid the formation of multiple channels.

**NaPSS and PLL:** The polyanion used in this work, sodium polystyrene supfonate (NaPSS) was purchased from Scientific Polymer Products NY, with weight-averaged molecular weights $M_w$= 1.5, 5.18, 16, 33.4, 126.7, and 262.8 kD. The polydispersity indices are 1.12, 1.13, 1.13, 1.17, 1.17 and 1.20, respectively. The polycation used, poly-L-lysine (PLL) was purchased from Alamanda polymers with average molecular weights of 1.6 ($PLL_{10}$), 3.3 ($PLL_{20}$), and 4.9 kD ($PLL_{30}$). After the insertion of a single αHL pore from the *cis* compartment, the solutions of *cis*/*trans* chambers were replaced with the KCl buffer solutions containing desired concentration of NaPSS and PLL respectively.

**ssDNA:** The sequence of 92-nt DNA was selected to have minimum secondary structure (1). 5' - AAAAAAAAAAAAAAAAAAAAATTCCCCCCCCCCCCCCCCCCCCCTTAAAAAAAAAATTCCCCCCCCCC TTAAAAAAAAAATTCCCCCCCCCC-3'**.** The sample of ssDNA was purchased from integrated DNA technology via custom oligo services (IDT USA). After the formation of single αHL pore, the t*rans* chambers were replaced with the KCl buffer solutions containing desired concentration of poly-lysine and then 50 μl of 15-20 μM of ssDNA were added to *cis* side.

**Electrical measurements and data analysis:** Ionic currents were measured by using Ag/AgCl electrodes with a patch-clamp amplifier (Axopatch 200B, Axon Instruments). The amplified signal was low-pass-filtered at 5 kHz (or 10 kHz) and sampled at 250 kHz with a Digidata 1550 digitizer (Axon Instruments). The recorded ionic current traces are analyzed using MATLAB (The MathWorks, Inc., MA) to capture the current blockage events and their durations.

**Analytical Model:** The model described was formulated and solved using *Mathematica*. The mathematical expression for $\tau_{cmp}$ was generated and then used to find the best fit to the entire data ($\tau_{cmp}$ vs voltage) set in the *Mathematica* software.

### (ii) Control experiments: NaPSS only translocation

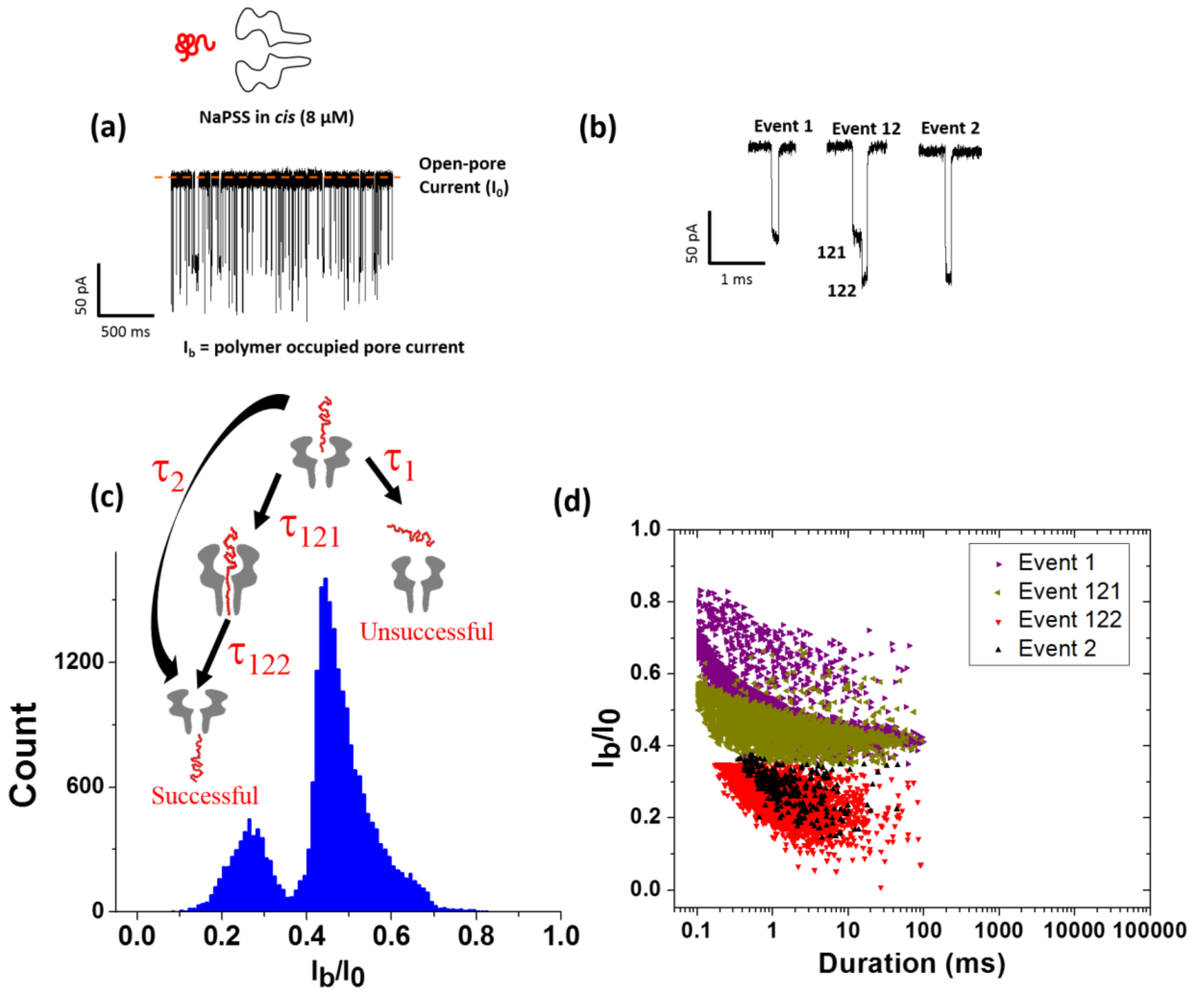


**Figure S1.** (a) Representative ionic current trace at 140mV, 7.5 pH, 30[0]C in 1M KCl, when 8 µM NaPSS(16kD) were present on *cis* compartment (b) Close look of different type of events in ionic current trace (c) Histogram of current blockade levels observed and (d) event diagram for NaPSS translocation at same condition. Histogram and event diagram shows two distinct population of observed events, one corresponds to successful ($I_b/I_0$ <0.35) and other is unsuccessful translocation ($I_b/I_0$ >0.35). Event type 1 and 121 are due to presence of NaPSS in vestibule.

### Control experiments: NaPSS only translocation

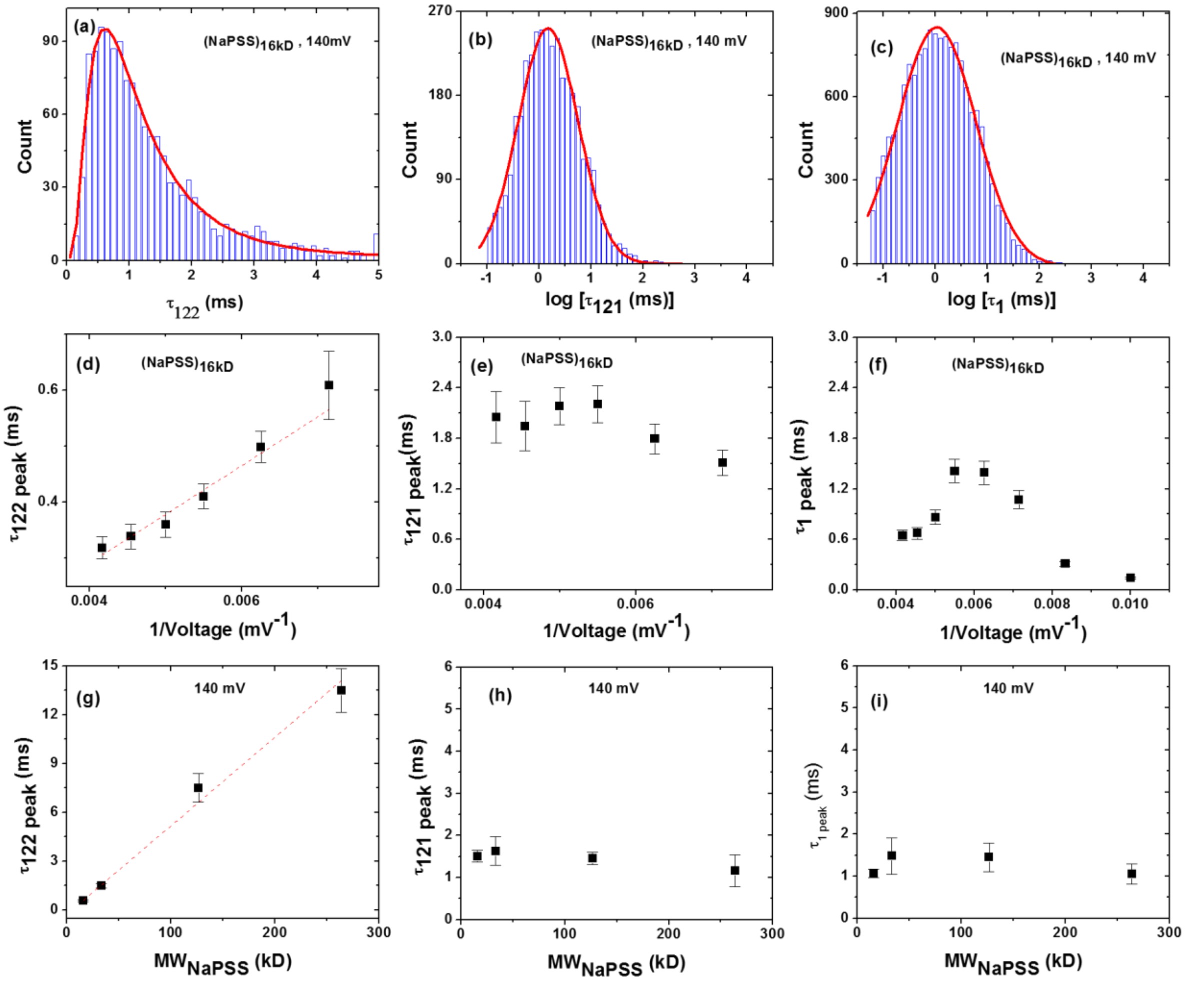

**Figure S2.** Histogram of duration of (a) Event122 (b) Event121 (c) Event1 for 16kD NaPSS at 140 mV, 7.5pH, 30$^0$C, in 1M KCl. Most probable duration as a function of voltage (d) for Event122 (e) for Event 121 (f) for Event1. Most probable duration as a function of molecular weight of NaPSS at 140 mV (g) for Event122 (h) for Event 121 (i) for Event1. Since, the populations of Event2 were very few they were analyzed together with Event 122. Error bars represent standard deviation from three independent experiments.

The durations of events of type 122 and 2 are proportional to molecular weight of NaPSS and inversely proportional to voltage. Therefore, only type122 and type2 of events are successful translocation events.

### Control experiments: PLL only translocation

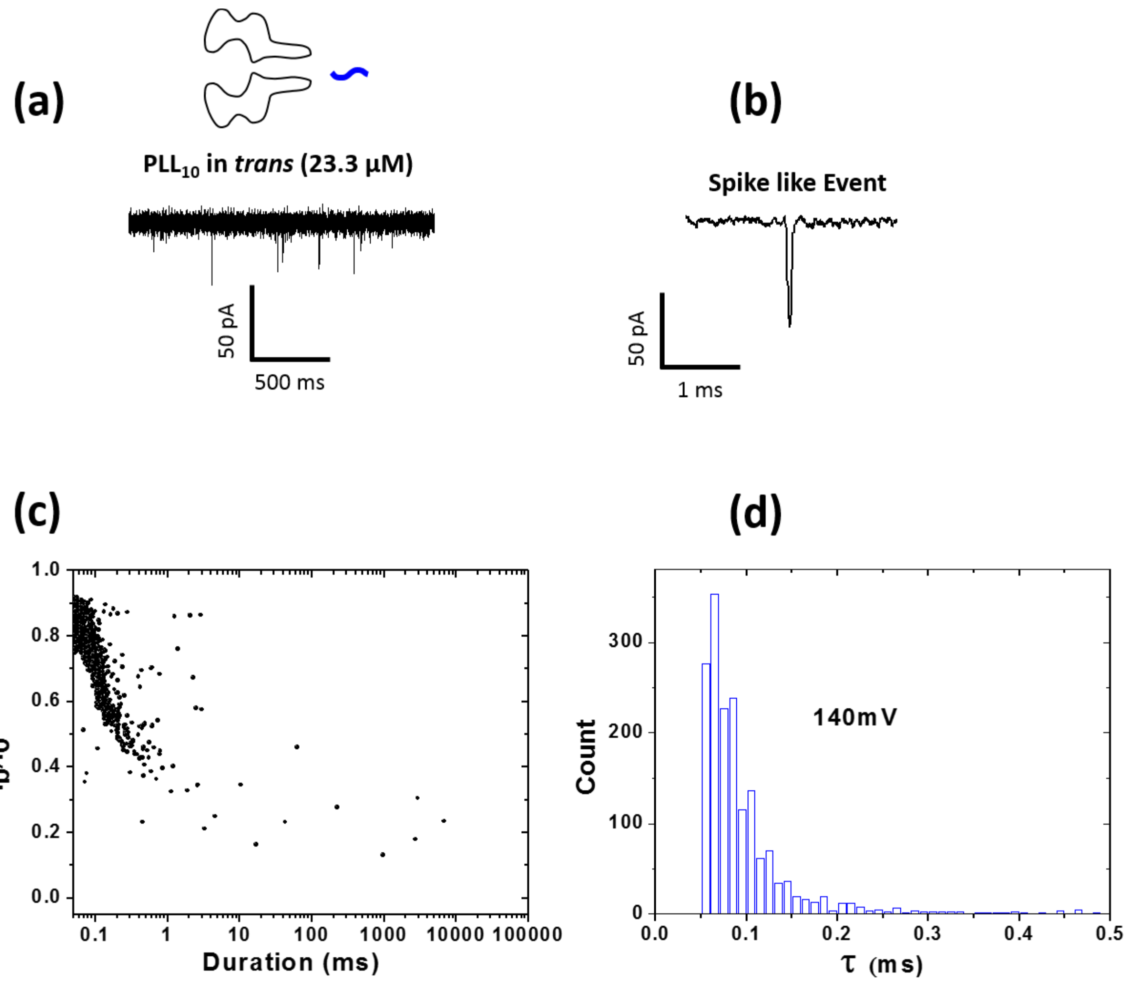


**Figure S3.** (a) Representative ionic current trace at 140mV, 7.5 pH, 30$^0$C in 1M KCl , when 23.3 μM PLL (1.6kD) were present on *trans* compartment (b) close look of events, mostly spike like event observed.(c) Event diagram showing depth of blocked pore current and its duration. (d) Histogram of the dwell time of PLL.

**Control experiments: NaPSS and PLL capture rates**

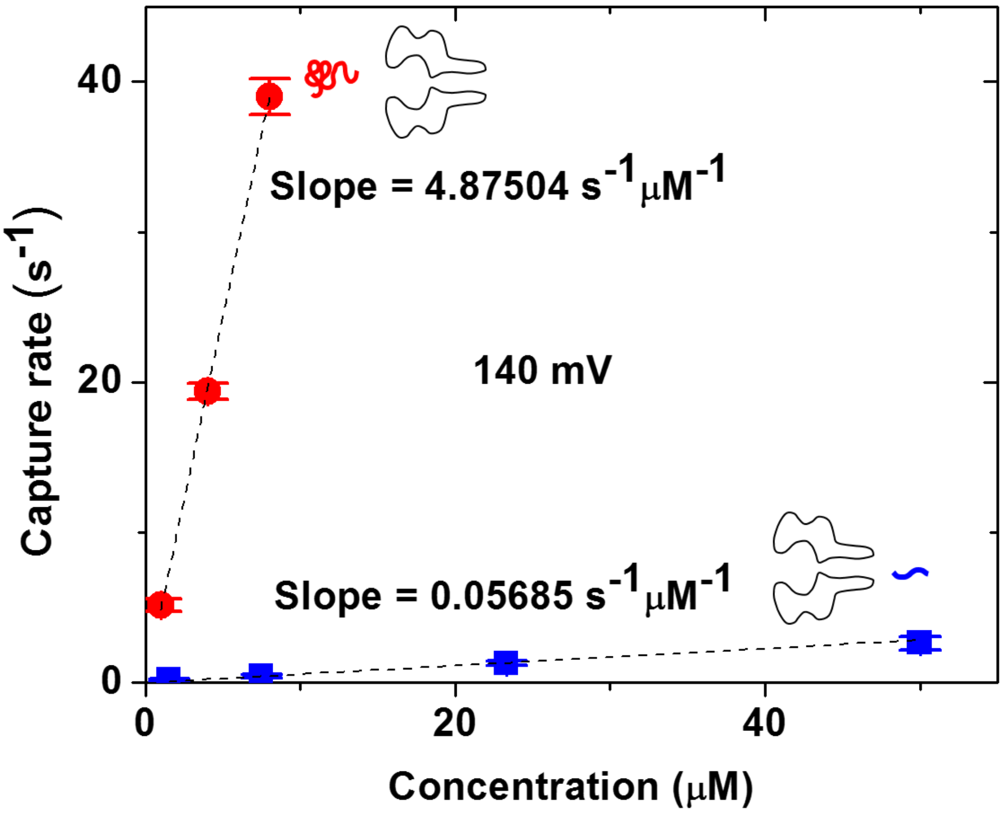


**Figure S4** Capture rate of NaPSS (16 kD) and PLL(1.6 kD) as a function of their concentration in *cis* and *trans* compartment respectively, at 140 mV, 7.5pH, 30$^0$C, in 1M KCl. Error bars represent standard deviation from three independent experiments.

Since, capture of NaPSS is 85 times more frequent compare to capture of PLL, therefore the probability that PLL will find an unoccupied pore is least.

### (iii) Additional complexation data

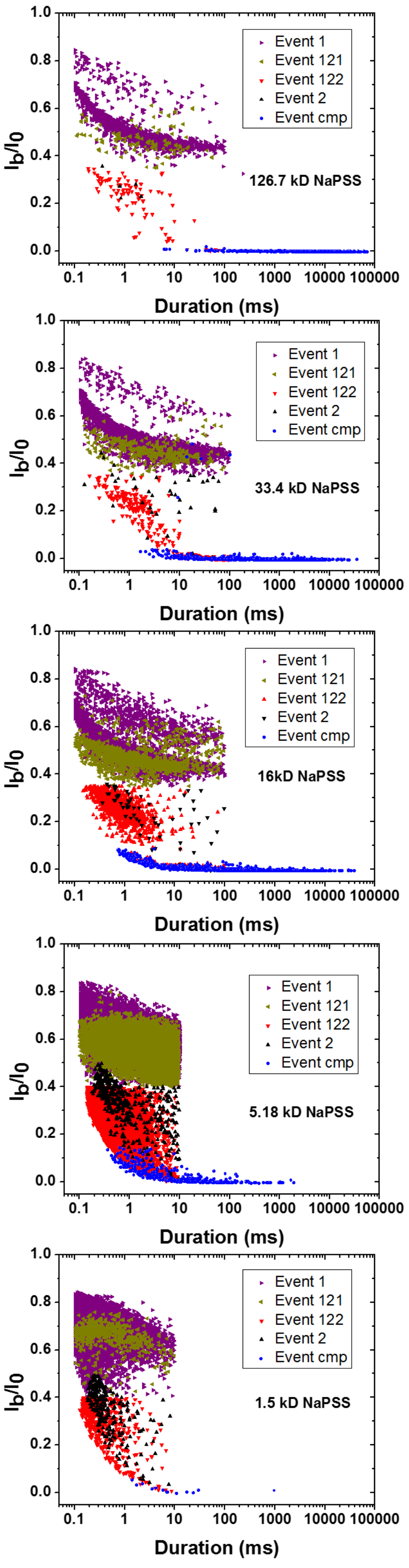


**Figure S5.** Event diagram of current blockade level when 8μM of different molecular weight of NaPSS were taken on *cis* side and 23.3 μM of $PLL_{1.6kD}$ were taken on *trans* side at 140 mV, 7.5 pH, $30^0$C in 1M KCl.

We can see from the event diagrams that, for higher molecular weight of NaPSS there are more population of complexation event. However, when the molecular weight of NaPSS decreases, population of complexation event also decreases. This is because longer NaPSS stay for longer time in the β-barrel and chances of its complexation with PLL increases.

### Additional complexation data

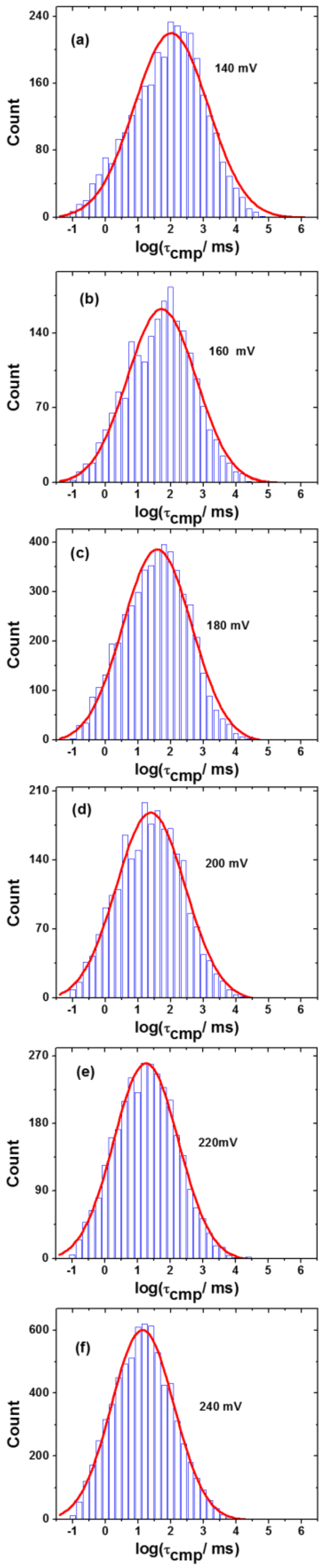


**Figure S6.** Histogram of log of complexation event duration at different voltages, NaPSS(16kD) and (23.3 μM) $PLL_{10}$. The histogram is fitted with log-normal probability density function.

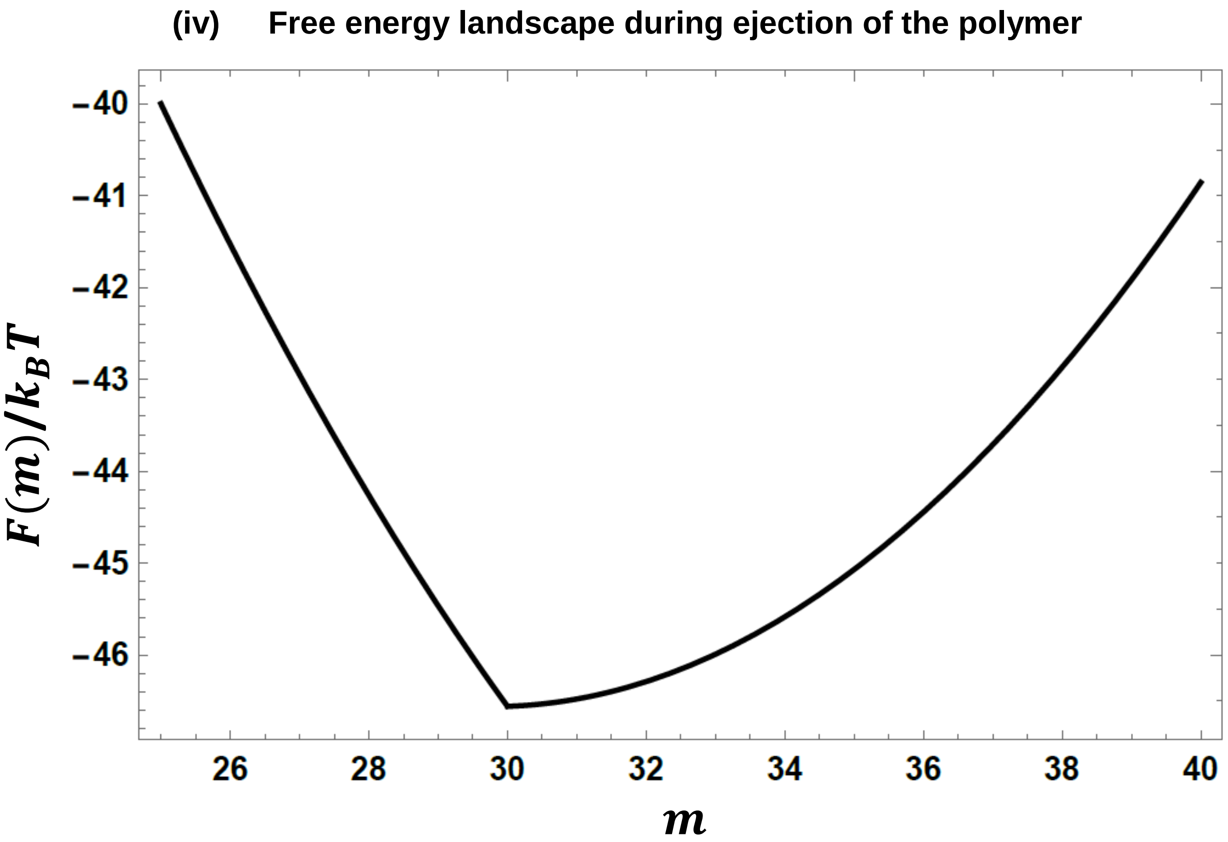


**Figure S7.** Free energy landscape during the ejection of the polymer (5.18kD NaPSS) according to the model described in main text ($\varepsilon_1 = 0.05$, $\varepsilon_2 = 1.115$, q=0.30, $N_1$=25, $N_2$=10). Energy barrier ($\nabla f$) is 5.702 $K_B T$.

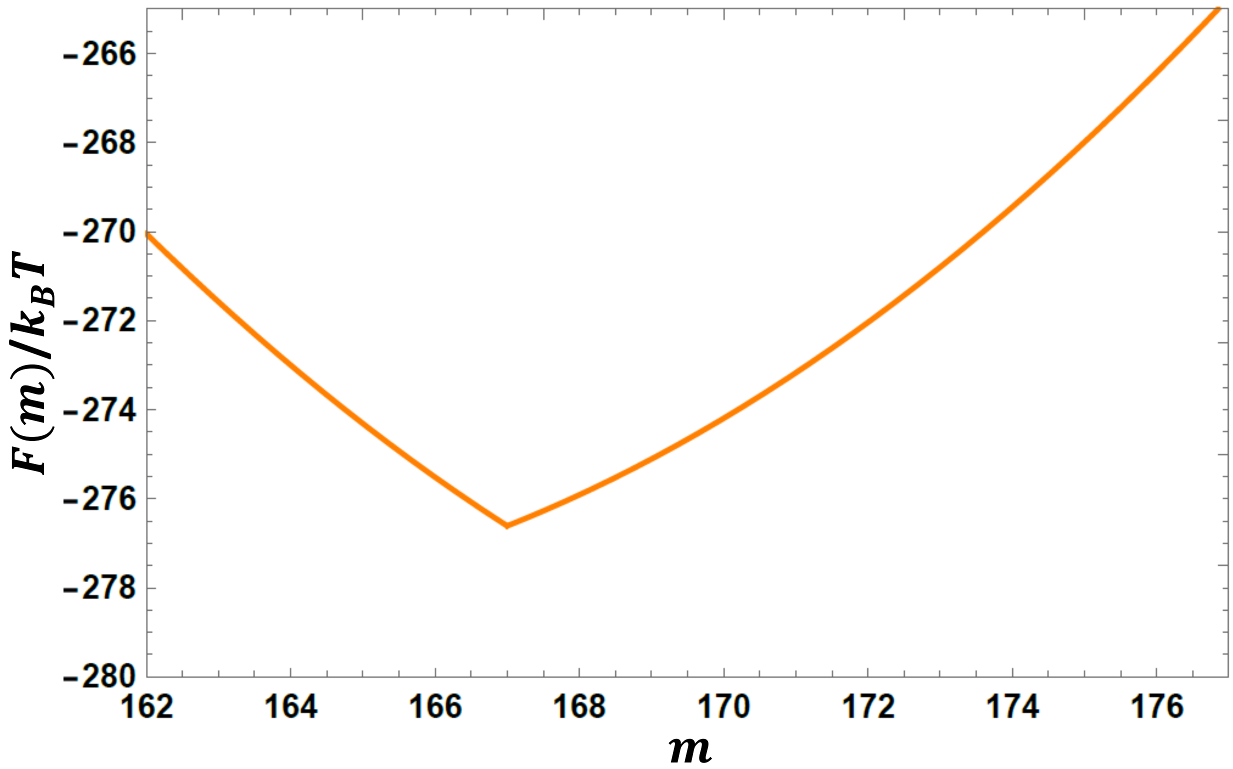


**Figure S8.** Free energy landscape during the ejection of the polymer (33.4kD NaPSS) according to the model described in main text ($\varepsilon_1 = 0.05$, $\varepsilon_2 = 1.731$, q=0.30, $N_1$=162, $N_2$=10). Energy barrier ($\nabla f$) is 11.862 $K_BT$.

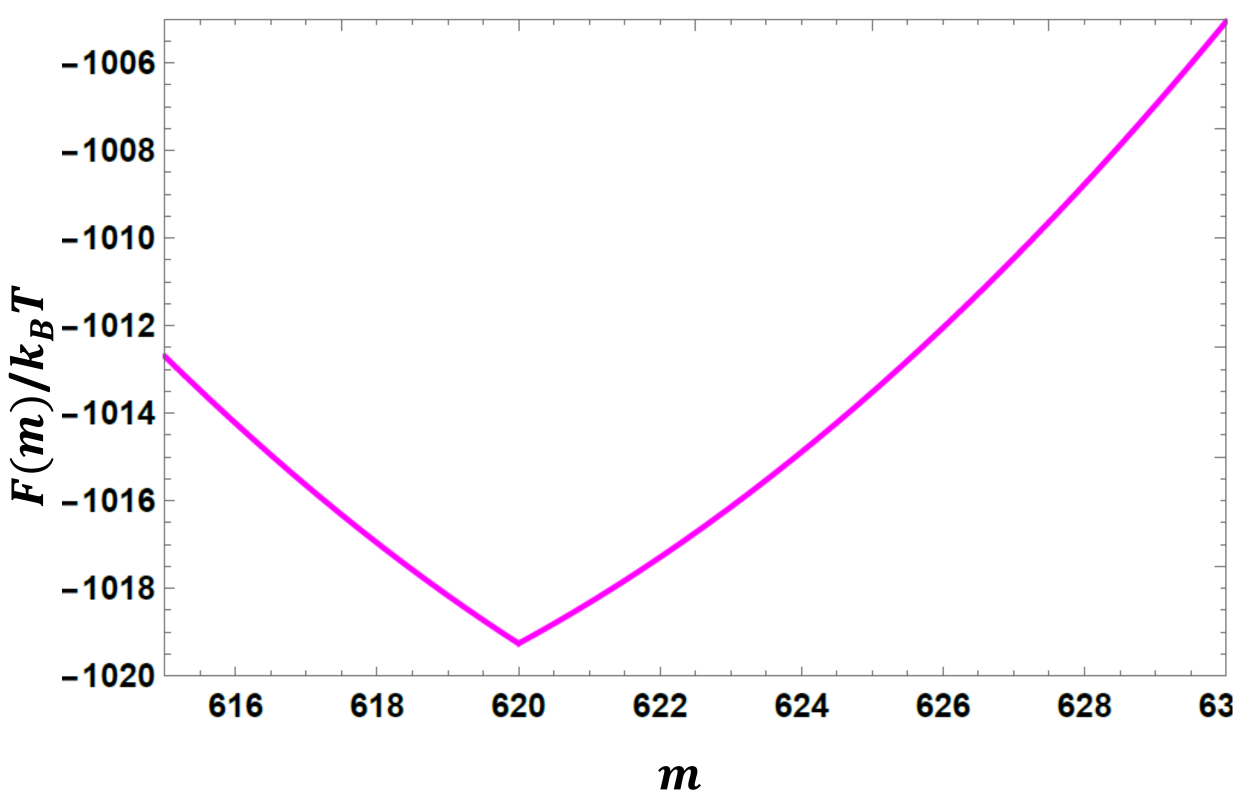


**Figure S9.** Free energy landscape during the ejection of the polymer (126.7kD NaPSS) according to the model described in main text ($\varepsilon_1 = 0.05$, $\varepsilon_2 = 1.965$, q=0.30, $N_1$=615, $N_2$=10). Energy barrier ($\nabla f$) is 14.202 $K_BT$.

### (v) Fit of the experimental data with the model

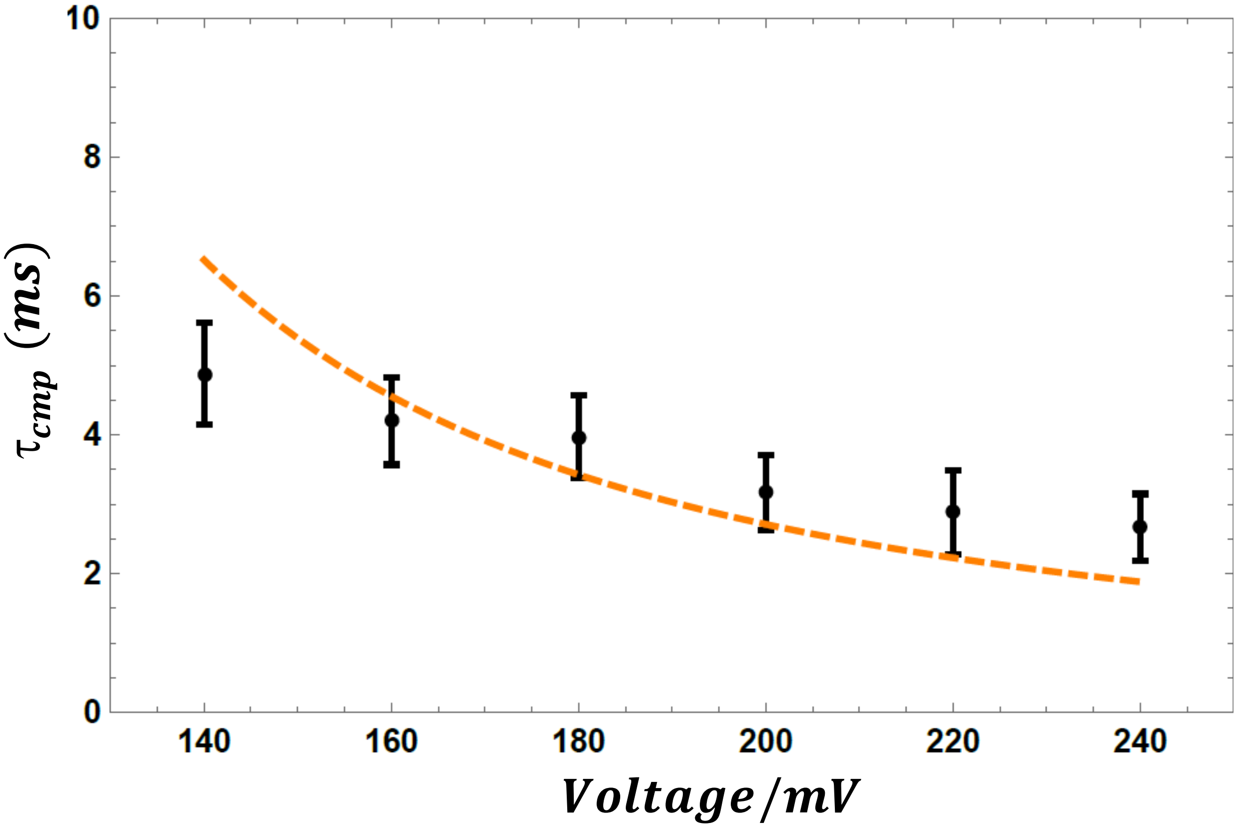


**Figure S10.** Experimentally observed and calculated values of $\tau_{cmp}$ are plotted together. The black dot with error bars represents experimental data for the 5.18 kD NaPSS and 1.6kD PLL. The orange dash curve represents fit from the model described in main text ($\varepsilon_1 = 0.05$, $\varepsilon_2 = 1.115$, q=0.30, $N_1$=25, $N_2$=10).

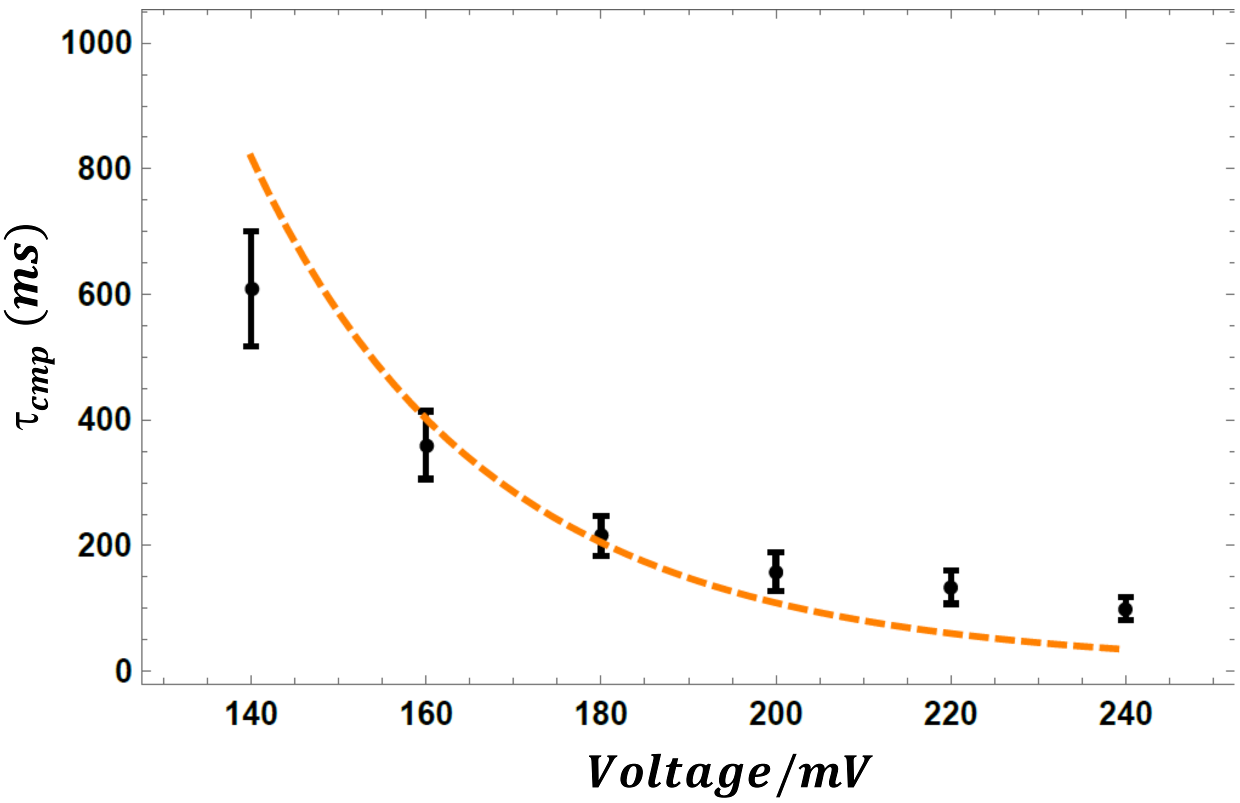


**Figure S11.** Experimentally observed and calculated values of $\tau_{cmp}$ are plotted together. The black dot with error bars represents experimental data for the 33.4 kD NaPSS and 1.6kD PLL. The orange dash curve represents fit from the model described in the main text ($\varepsilon_1 = 0.05$, $\varepsilon_2 = 1.731$, q=0.30, $N_1$=162, $N_2$=10).

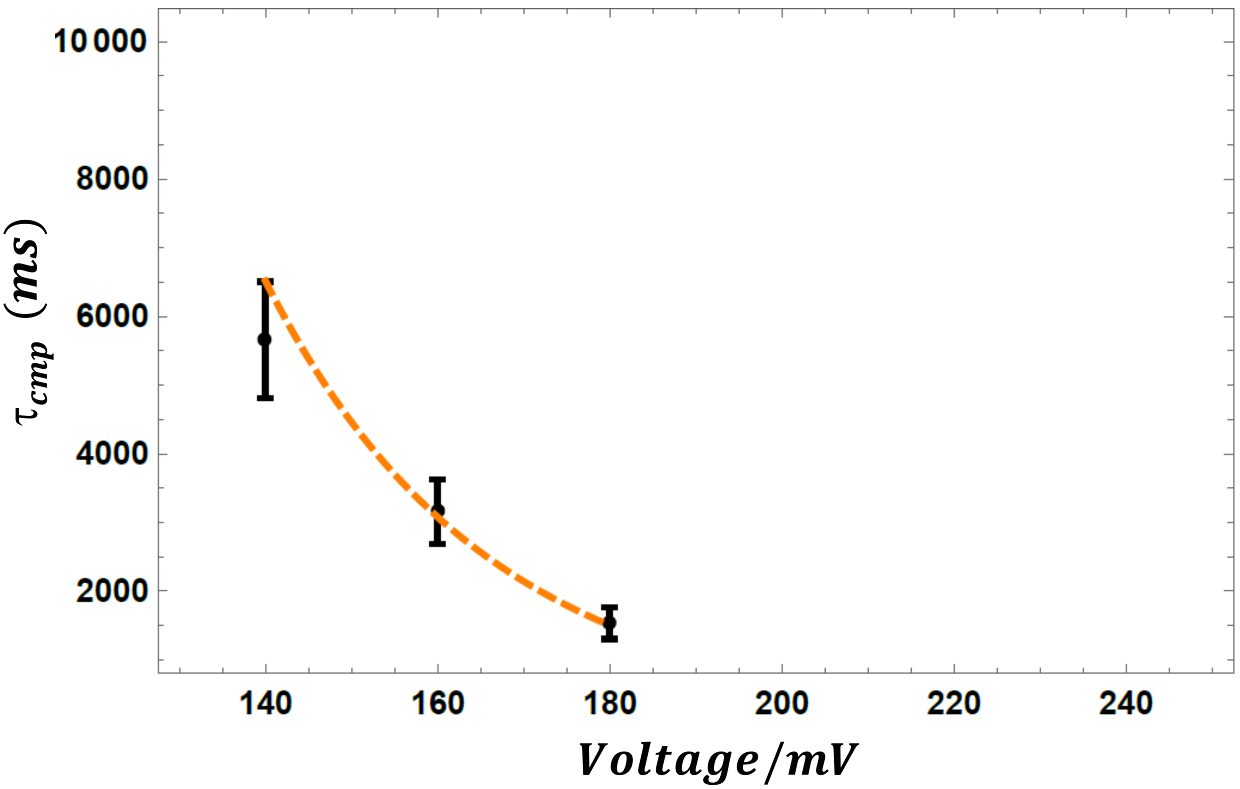


**Figure S12.** Experimentally observed and calculated values of $\tau_{cmp}$ are plotted together. The black dot with error bars represents experimental data for the 126.7 kD NaPSS and 1.6kD PLL. The orange dash curve represents fit from the model described in the main text ($\varepsilon_1 = 0.05$, $\varepsilon_2 = 1.965$, q=0.30, $N_1$=615, $N_2$=10).

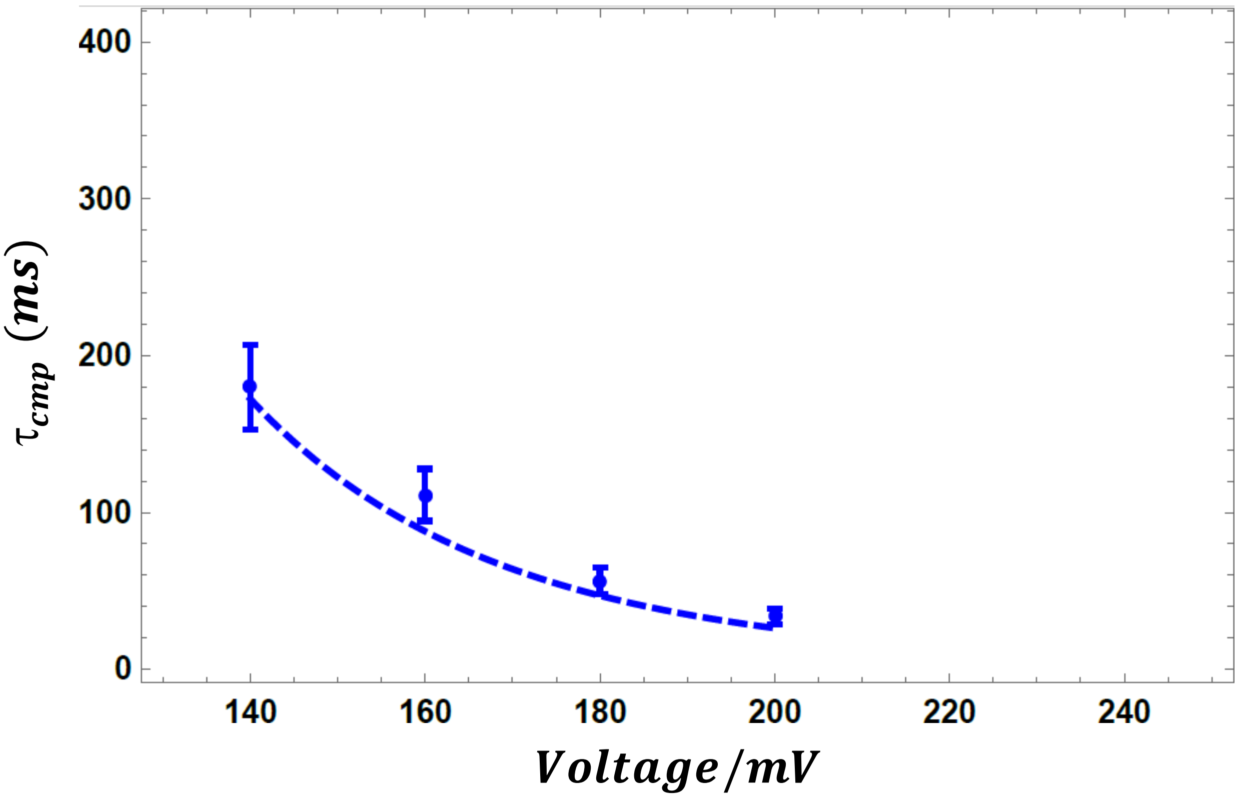


**Figure S13.** Experimentally observed and calculated values of $\tau_{cmp}$ are plotted together. The blue dot with error bars represents experimental data for the 16 kD NaPSS and 3.3kD PLL. The blue dash curve represents fit from the model described in the main text ($\varepsilon_1 = 0.05$, $\varepsilon_2 = 1.550$, q=0.30, $N_1$=78, $N_2$=10).

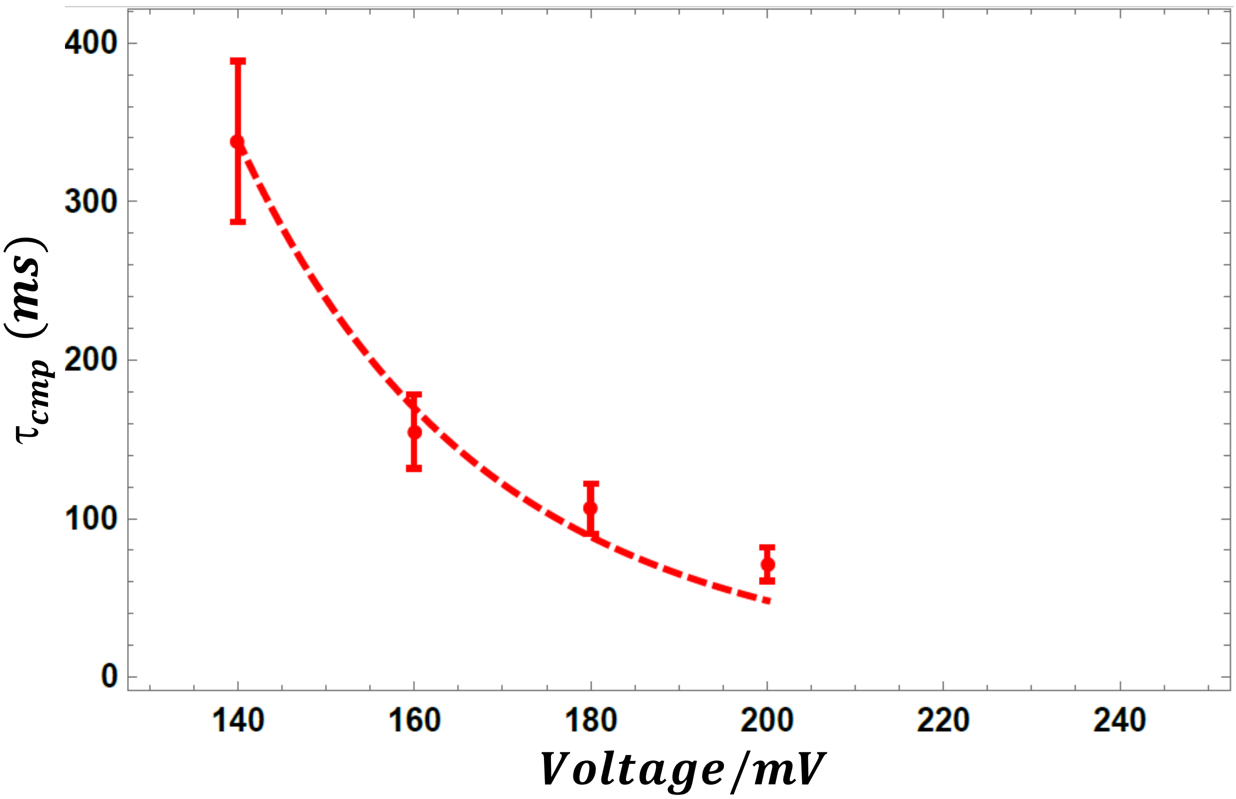


**Figure S14.** Experimentally observed and calculated values of $\tau_{cmp}$ are plotted together. The red dot with error bars represents experimental data for the 16 kD NaPSS and 4.9kD PLL. The red dash curve represents fit from the model described in the main text ($\varepsilon_1 = 0.05$, $\varepsilon_2 = 1.630$, q=0.30, $N_1$=78, $N_2$=10).

#### (vi) Physical basis for double-exponential forms

Given that the complex is already inside the pore at time t, the probability of not finding the complex inside the pore in a time interval t to t+Δt, is proportional to Δt at first order. Therefore, the probability of finding the complex inside the pore in a time interval t to t+Δt is:

$$p(t \rightarrow \mathrm{t} + \Delta\,\mathrm{t}) = 1 - \alpha\,\Delta\,\mathrm{t}\,; \quad (1)$$

where, $\alpha$ is a constant independent of time.

Let the probability density of finding a complex with dwell time t be p(t). The probability density p(t+Δt) of the complex having dwell time t+Δt will be proportional to:

(i) The probability density p(t) of finding the complex with dwell time t.

(ii) The probability $p(t \rightarrow \mathrm{t} + \Delta\mathrm{t})$ of finding the complex inside the pore in the following time interval, t to t+Δt.

The above two events are mutually independent, therefore:

$$\mathrm{p}(\mathrm{t} + \Delta\mathrm{t}) = \mathrm{p}(\mathrm{t})(1 - \alpha\,\Delta\mathrm{t}\,); \quad (2)$$

$$\mathrm{p}(\mathrm{t} + \Delta\mathrm{t}) - \mathrm{p}(\mathrm{t}) = -\,\alpha\,\Delta\mathrm{t}\,\mathrm{p}(\mathrm{t})\,; \quad (3)$$

$$\frac{\mathrm{p}(\mathrm{t} + \Delta\mathrm{t}) - \mathrm{p}(\mathrm{t})}{\Delta\mathrm{t}} = -\,\alpha\,\mathrm{p}(\mathrm{t})\,; \quad (4)$$

By definition of the derivative, in the limit of Δt → 0, the above equation becomes:

$$\frac{\mathrm{d}\,\mathrm{p}(\mathrm{t})}{\mathrm{dt}} = -\,\alpha\,\mathrm{p}(\mathrm{t})\,; \quad (5)$$

and by integration we obtain:

$$\mathrm{p}(\mathrm{t}) = \mathrm{a}\,e^{-\,\alpha\,t}; \quad (6)$$

The quantity $\alpha$ has a unit of inverse of time can be written as $1/\tau$, where $\tau$ is characteristic time constant for the complex life time distribution. The constant a is determined by normalization to be $a = \alpha$.

However, the histogram of complex life-time gives a poor fit with single exponential probability density function. Therefore, we implement double exponential forms and we describe probability density function for the complex life-time having two-time constants, given by

$$f(t) = \frac{a_1}{\tau_1}\, e^{-\frac{t}{\tau_1}} + \frac{a_2}{\tau_2}\, e^{-\frac{t}{\tau_2}} \;; \qquad (7)$$

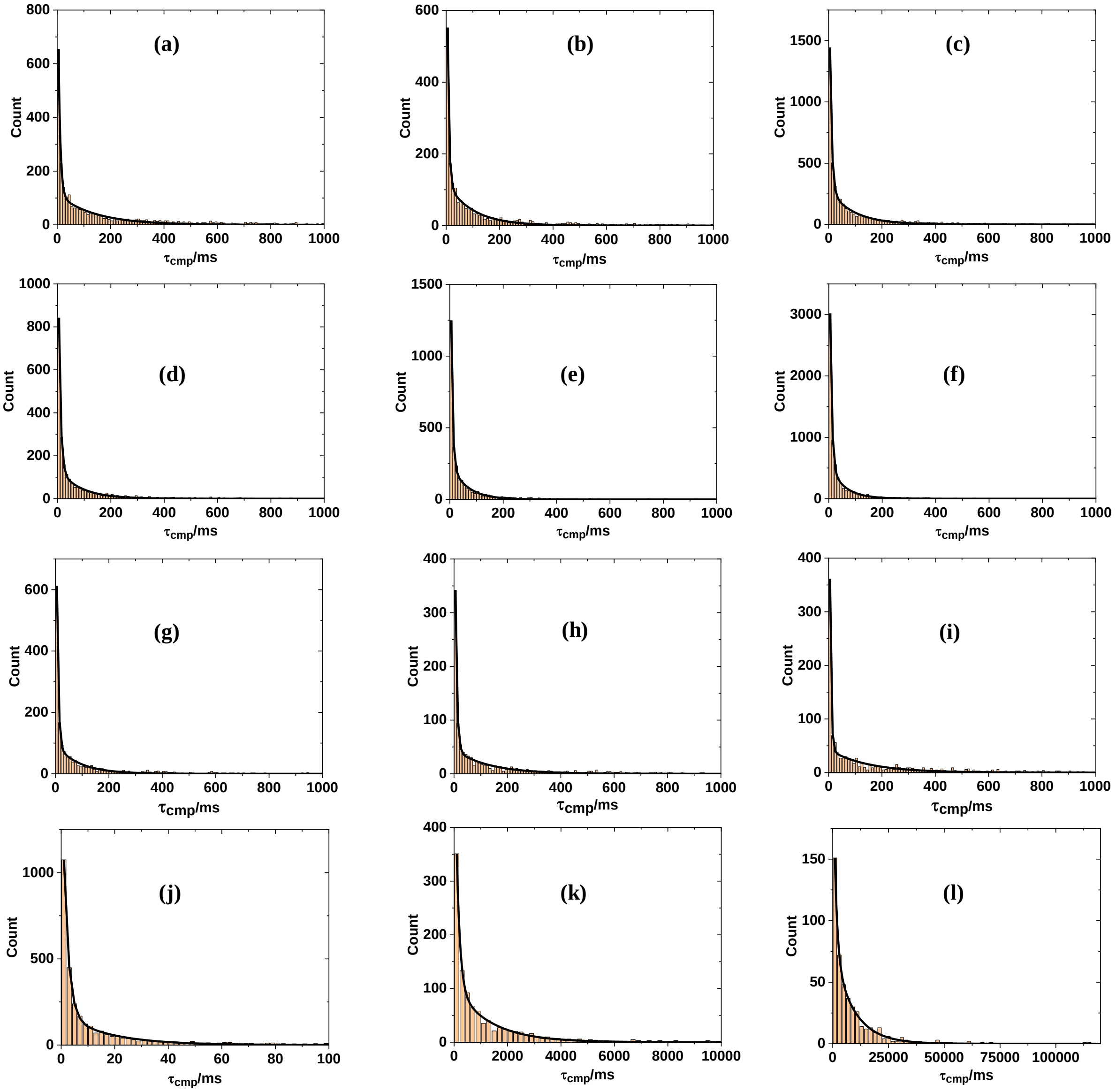


**Figure S15**. Histogram of the life-time of the complex for 16kD PSS and 1.6kD PLL 23.3μM at **(a)** 140mV for 3079 events **(b)** 160mV for 2104 events **(c)** 180mV for 4955 events **(d)** 200mV for 2413 events **(e)** 220mV for 3144 events **(f)** 240mV for 7036 events, At 140mV different concertation of PLL **(g)** 1.5 μM PLL 1809 events **(h)** 7.5 μM PLL for 1170 events **(i)** 50 μM PLL for 1251 events, and for different molecular weight of NaPSS for **(j)** 5.18kD PSS at 140mV for 3416 events **(k)** 33.4 kD PSS 140mV 1065 events **(l)** 126.7 kD PSS at 140mV for 466 events. The black solid curve represents fit of the histogram with double exponential probability density function.

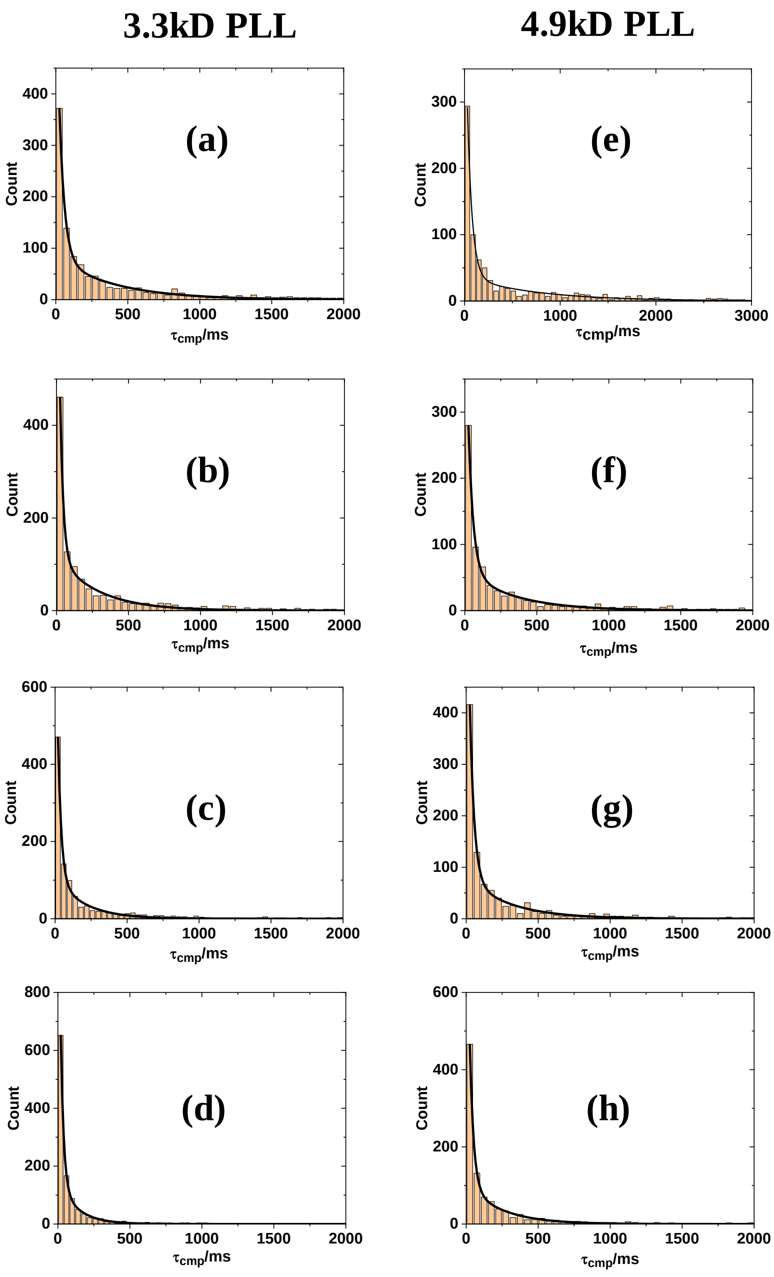


**Figure S16**. Histogram of the life-time of the complex for 16kD PSS and 3.3kD PLL (23.3μM) at **(a)** 140mV for 1195 events **(b)** 160mV for 1208 events **(c)** 180mV for 1116 events **(d)** 200mV for 1200 events, and for 4.9kD PLL (23.3 μM) at **(e)** 140mV for 920 events **(f)** 160mV for 801 events **(g)** 180mV for 1001 events **(h)** 200mV for 1026 events. The black solid curve represents fit of the histogram with double exponential probability density function.

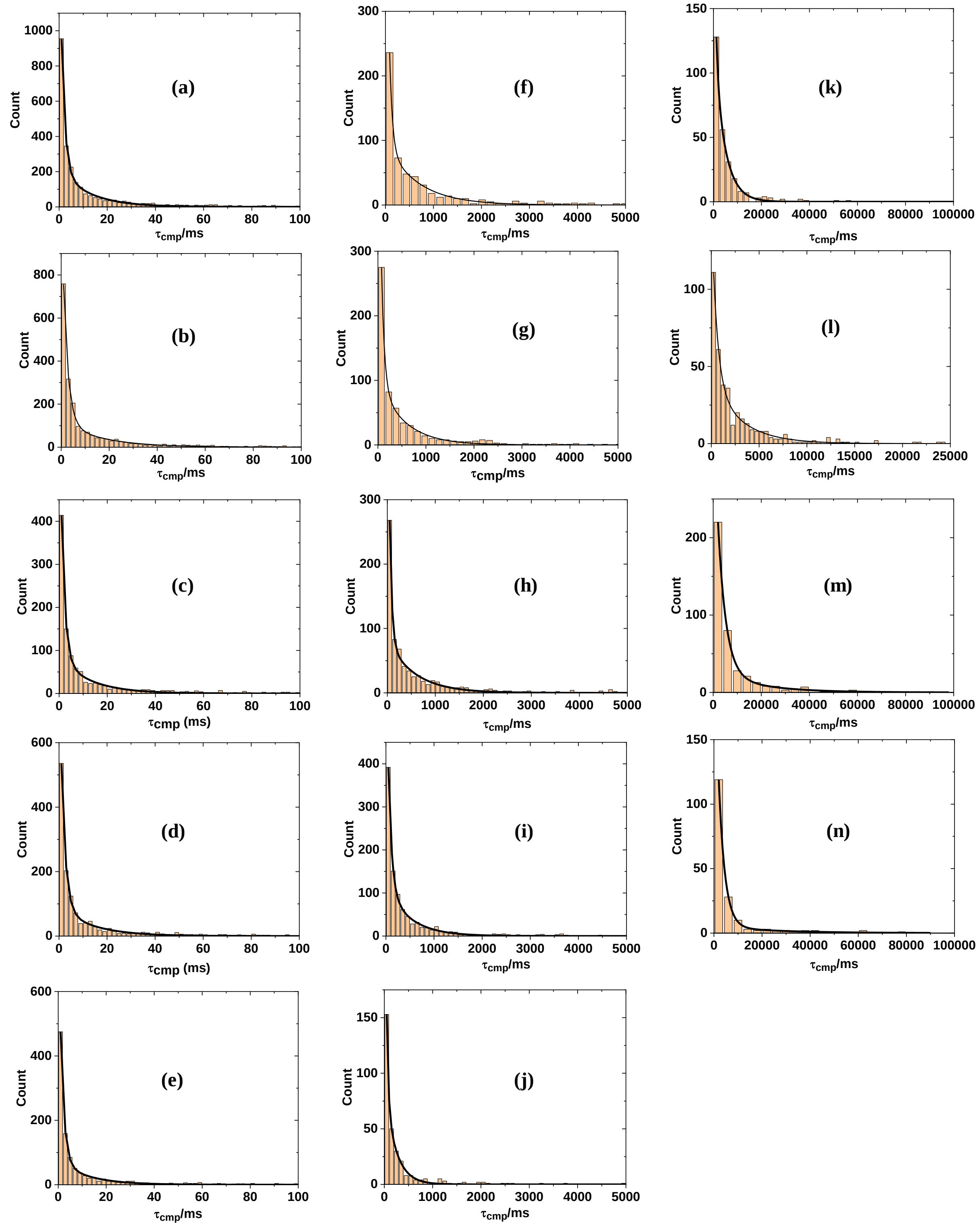


**Figure S17**. Histogram of the life-time of the complex for 5.18kD PSS and 1.6kD PLL (23.3μM) at (a) 160mV for 2783 events (b) 180mV for 2242 events (c) 200mV for 1185 events (d) 220mV for 1468 events (e) 240mV for 1153 events. For 33.4kD NaPSS and 1.6kD PLL (23.3μM) (f) 160mV for 579 events. (g) 180mV for 613 events (h) 200mV for 764 events (i) 220mV for 1045 events (j) 240mV for 316 events. For 126.7kD

NaPSS and 1.6kD PLL (23.3μM) (k) 160mV for 266 events (l) 180mV for 386 events (m) 200mV for 402 events (n) 220mV for 177 events

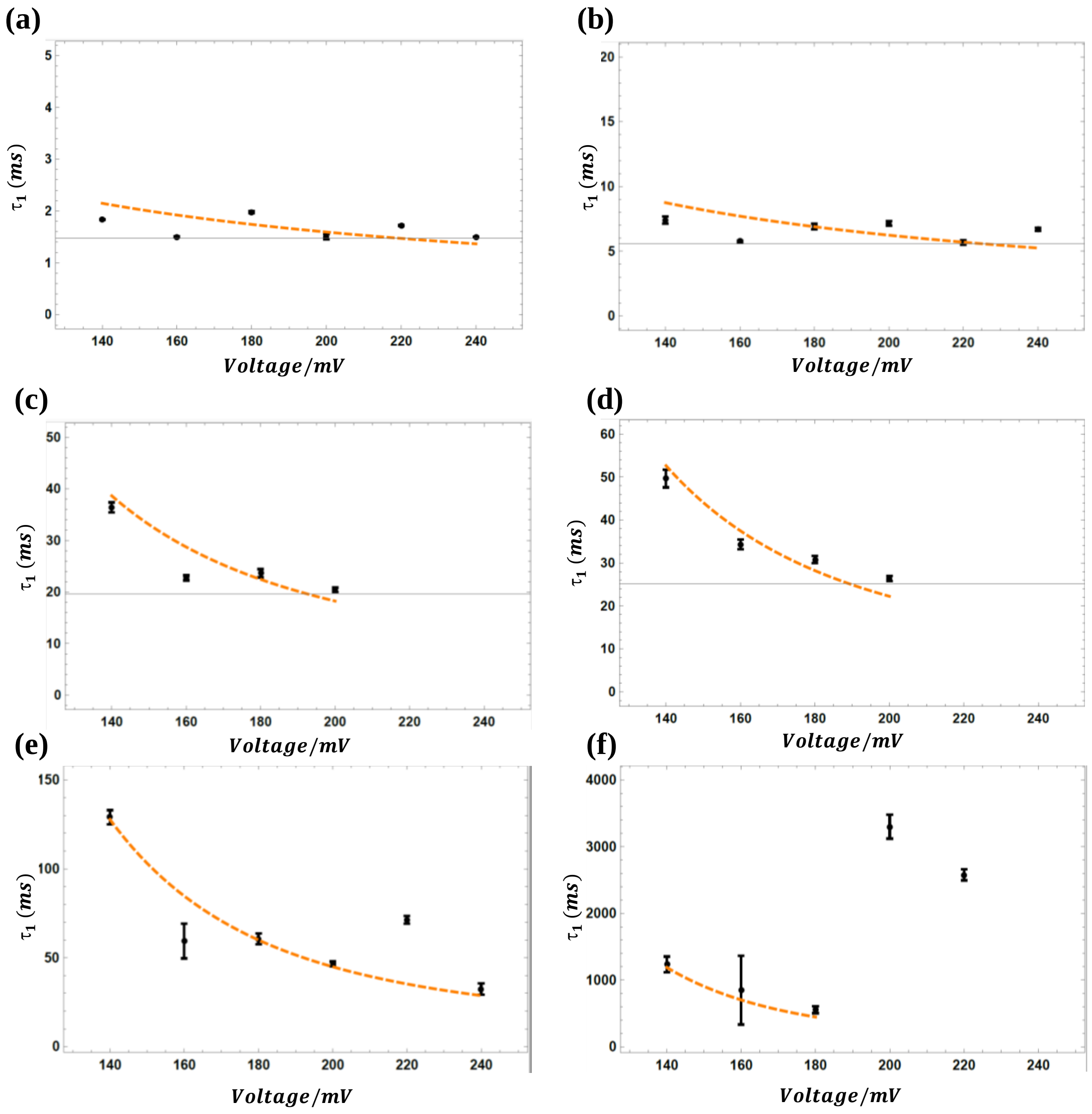


**Figure S18**. Experimentally observed and calculated values of $\tau_1$ are plotted together. (a) The black dot with error bars represents experimental data for the 5.18 kD NaPSS and 1.6 kD PLL (23.3 μM). The orange dash curve represents fit from the model described in section 2.6 ($\varepsilon 1 = 0.05$, $\varepsilon 2 = 0.01$, q=0.30, N1=25, N2=10). (b) 16 kD NaPSS and 1.6 kD PLL (23.3 μM). ($\varepsilon 1 = 0.05$, $\varepsilon 2 = 0.11$, q=0.30, N1=78, N2=10). (c)16 kD NaPSS and 3.3 kD PLL (23.3 μM). ($\varepsilon 1 = 0.05$, $\varepsilon 2 = 0.78$, q=0.30, N1=78, N2=10). (d) 16 kD NaPSS and 4.9 kD PLL (23.3 μM). ($\varepsilon 1 = 0.05$, $\varepsilon 2 = 0.85$, q=0.30, N1=78, N2=10). (e) 33.4 kD NaPSS and 1.6 kD PLL (23.3 μM). ($\varepsilon 1 = 0.05$, $\varepsilon 2 = 1.0$, q=0.30, N1=162, N2=10). (f) 126.7 kD NaPSS and 1.6 kD PLL (23.3

μM). ($\varepsilon 1 = 0.05$, $\varepsilon 2 = 1.25$, q=0.30, N1=615, N2=10). Error bars represents standard error obtained in fitting the histogram with double exponential probability density function.

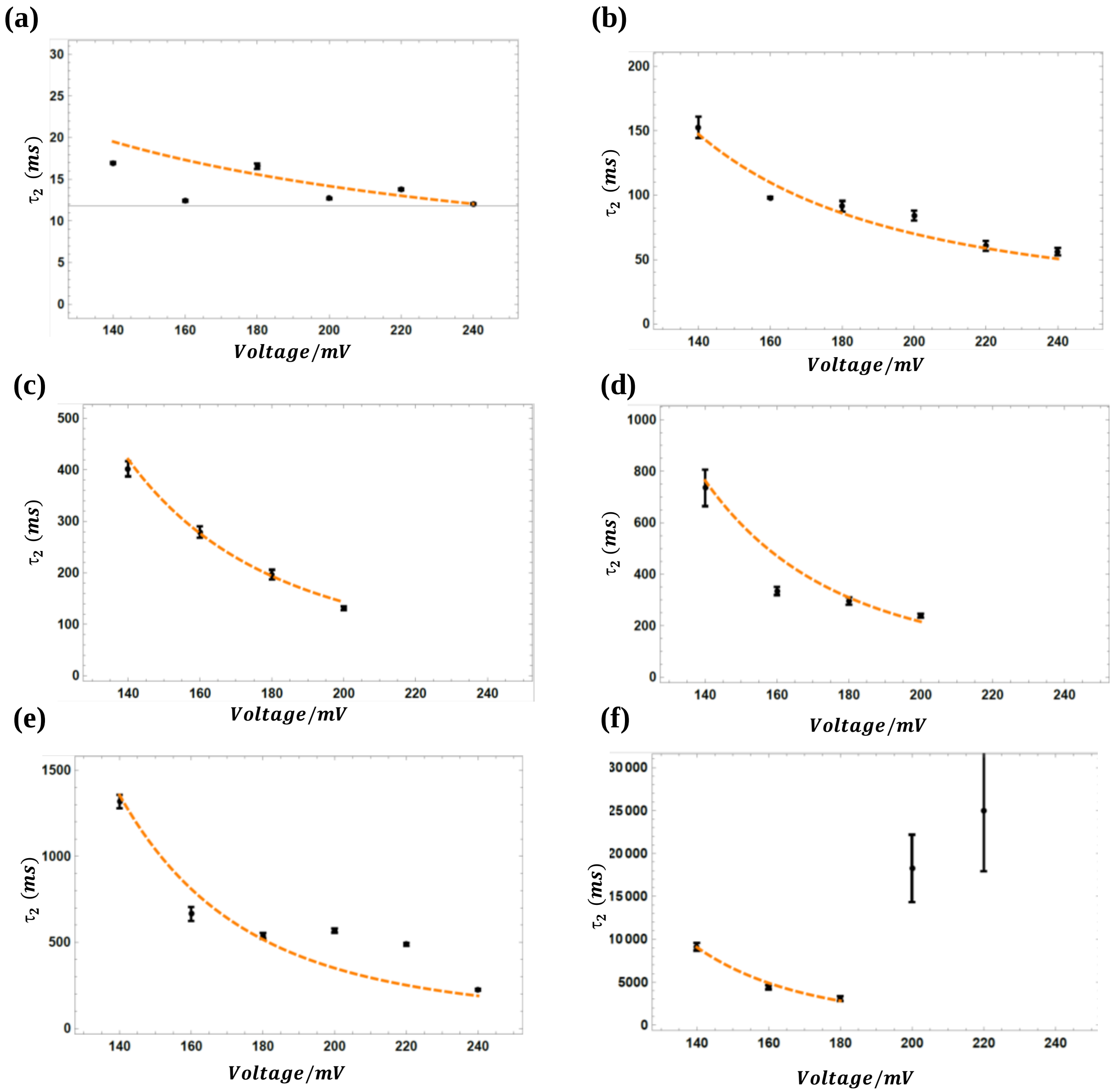


**Figure S19**. Experimentally observed and calculated values of $\tau_2$ are plotted together. (a) The black dot with error bars represents experimental data for the 5.18 kD NaPSS and 1.6 kD PLL (23.3 μM). The orange dash curve represents fit from the model described in section 2.6 ($\varepsilon 1 = 0.05$, $\varepsilon 2 = 0.15$, q=0.30, N1=25, N2=10). (b) 16 kD NaPSS and 1.6 kD PLL (23.3 μM). ($\varepsilon 1 = 0.05$, $\varepsilon 2 = 0.775$, q=0.30, N1=78, N2=10). (c)16 kD NaPSS and 3.3 kD PLL (23.3 μM). ($\varepsilon 1 = 0.05$, $\varepsilon 2 = 0.973$, q=0.30, N1=78, N2=10). (d) 16 kD NaPSS and 4.9 kD PLL (23.3 μM). ($\varepsilon 1 = 0.05$, $\varepsilon 2 = 1.08$, q=0.30, N1=78, N2=10). (e) 33.4 kD NaPSS and 1.6 kD PLL (23.3 μM). ($\varepsilon 1 = 0.05$, $\varepsilon 2 = 1.160$, q=0.30, N1=162, N2=10). (f) 126.7 kD NaPSS and 1.6 kD PLL (23.3 μM). ($\varepsilon 1 = 0.05$, $\varepsilon 2 = 1.41$, q=0.30, N1=615, N2=10). Error bars represents standard error obtained in fitting the histogram with double exponential probability density function

**S20: Table T1.** Values of interaction energy parameters $\varepsilon_2$ and free energy barrier $\Delta$f, obtained from best fit of calculated $\tau_1$ from the model described in main text and experimentally observed $\tau_1$.

| $N_{PLL}$ | $N_{PSS}$ | $\varepsilon_2\,(k_BT)$ | $\Delta f\,(k_BT)$ |
|---|---|---|---|
| 10 | 25 | 0.01 | 0.009 |
| 10 | 78 | 0.11 | 0.05 |
| 10 | 162 | 1.0 | 4.58 |
| 10 | 615 | 1.25 | 7.05 |
| 20 | 78 | 0.78 | 2.79 |
| 30 | 78 | 0.85 | 3.31 |

**S21: Table T2.** Values of interaction energy parameters $\varepsilon_2$ and free energy barrier $\Delta$f, obtained from best fit of calculated $\tau_2$ from the model described in main text and experimentally observed $\tau_2$.

| $N_{PLL}$ | $N_{PSS}$ | $\varepsilon_2\,(k_BT)$ | $\Delta f\,(k_BT)$ |
|---|---|---|---|
| 10 | 25 | 0.15 | 0.103 |
| 10 | 78 | 0.775 | 2.256 |
| 10 | 162 | 1.160 | 6.152 |
| 10 | 615 | 1.41 | 8.65 |
| 20 | 78 | 0.973 | 4.34 |
| 30 | 78 | 1.08 | 5.352 |

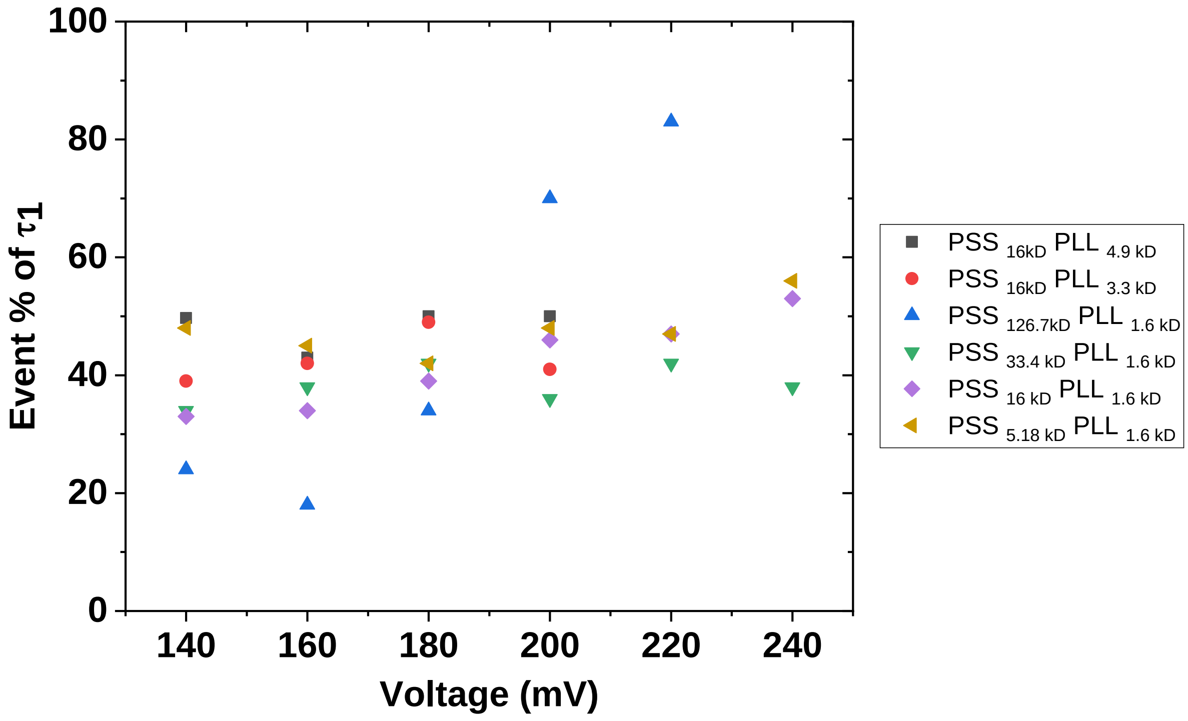


**S22:** Dependence of fraction of event with time constant $\tau_1$ on voltage, for all molecular weight of PSS and PLL.

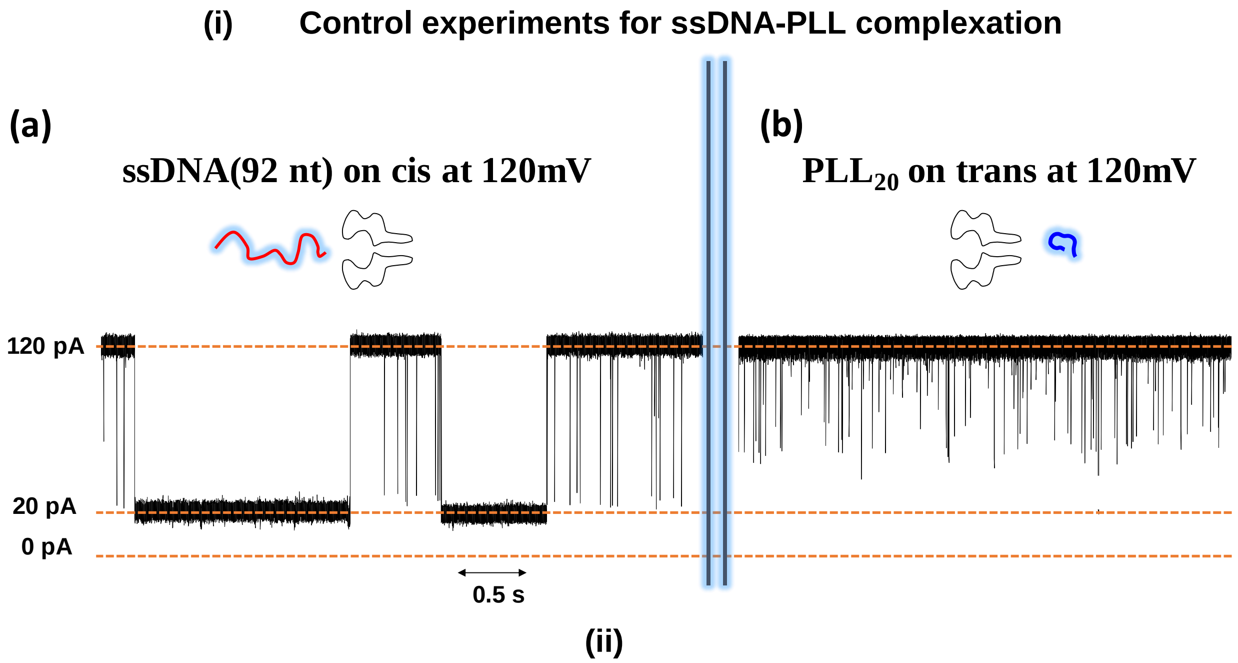


**Figure S23.** Representative ionic current trace at 120mV, 7.5pH, in 1M KCl, when **(a)** only 50 μl of 15-20 μM ssDNA was placed in *cis* compartment **(b)** only 50 μM of $PLL_{20}$ was placed in trans compartment.

**Control experiments**

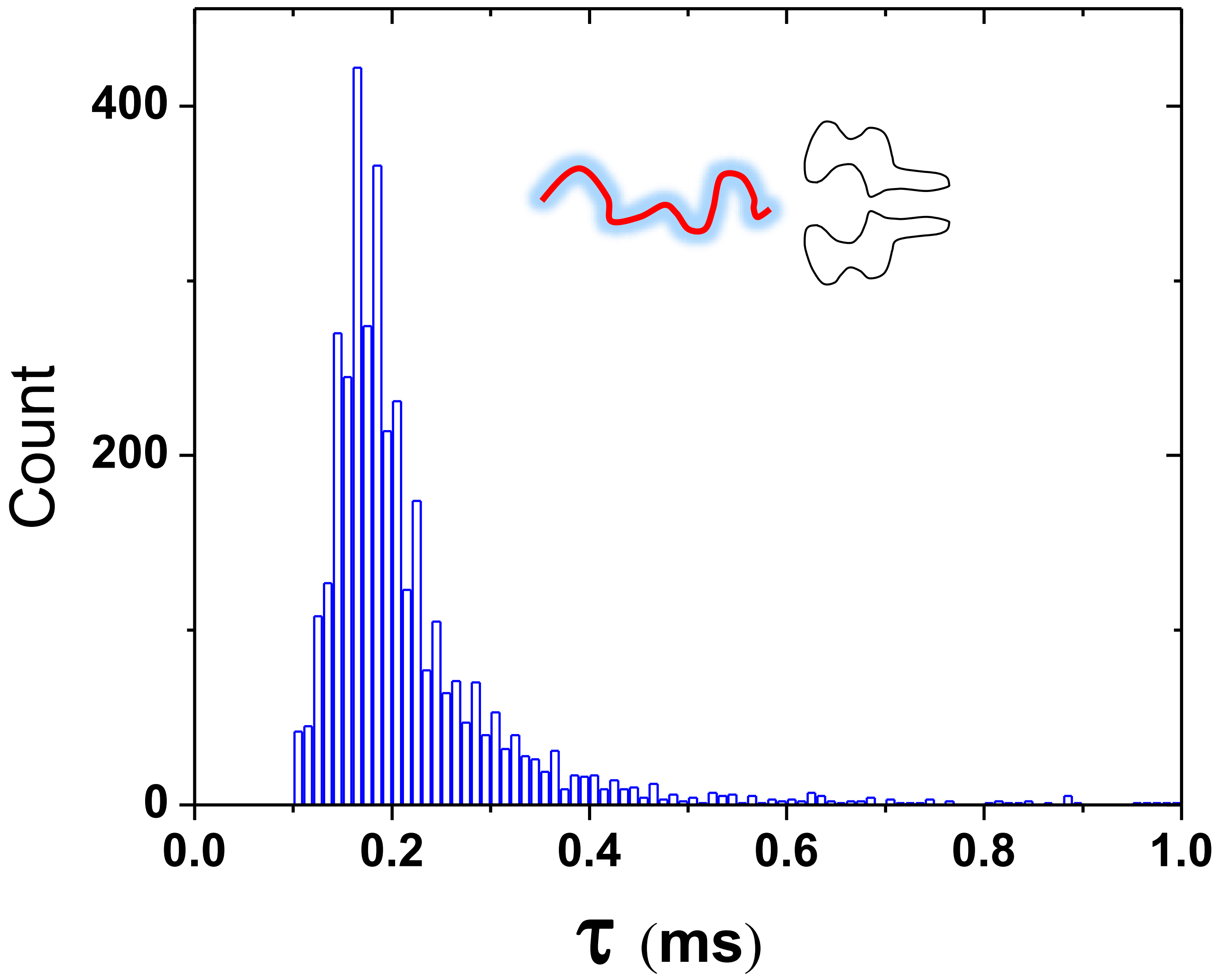


**Figure S24.** Histogram displaying dwell time of ssDNA during its translocation though the pore at 120 mV, 1M KCl and 7.5pH.

### Control experiments

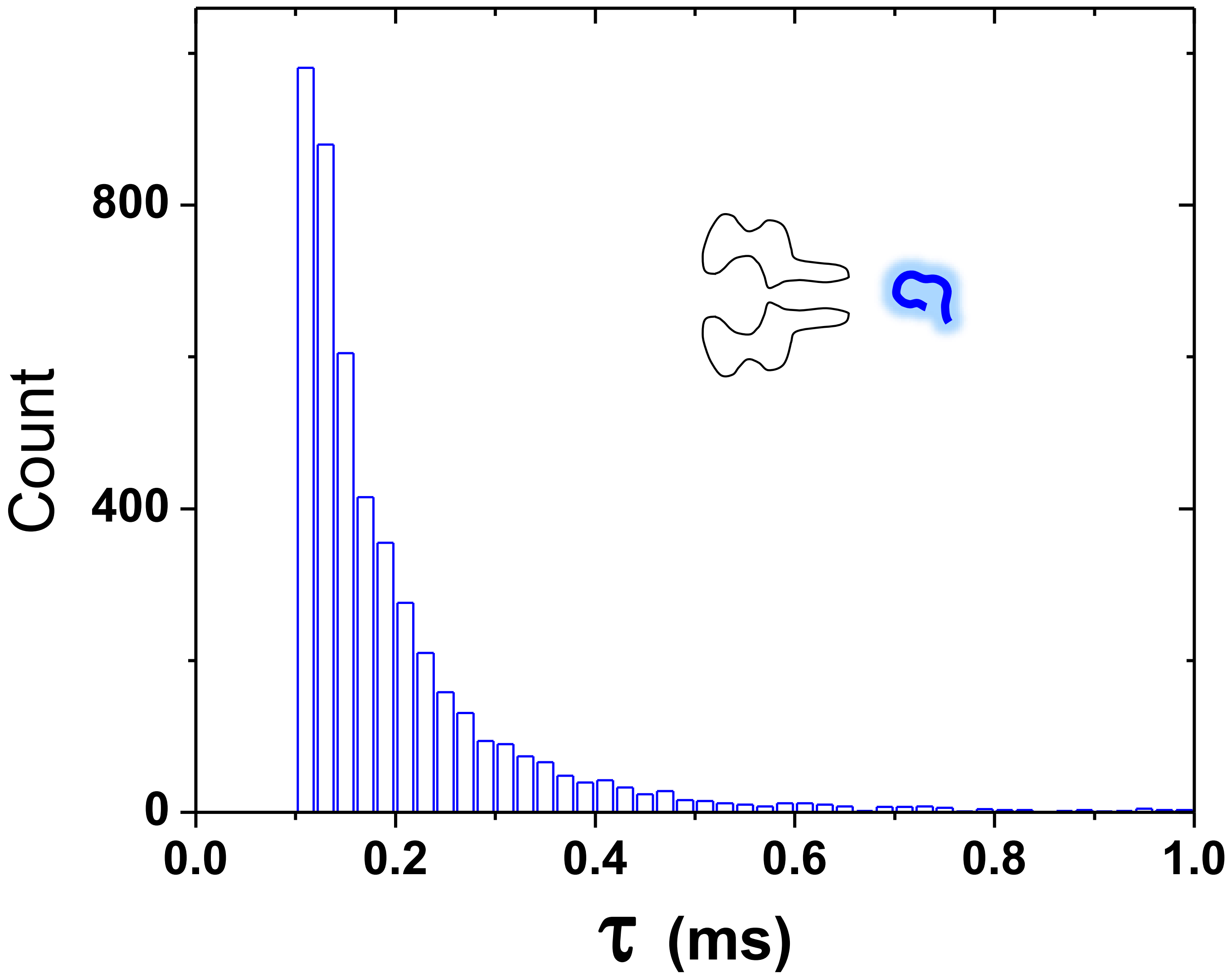


**Figure S25.** Histogram displaying dwell time of poly-L-lysine ($PLL_{20}$) for its interaction with the pore at 120 mV, 7.5 pH and 1M KCl.

### (iii) Additional data for ssDNA-PLL complexation

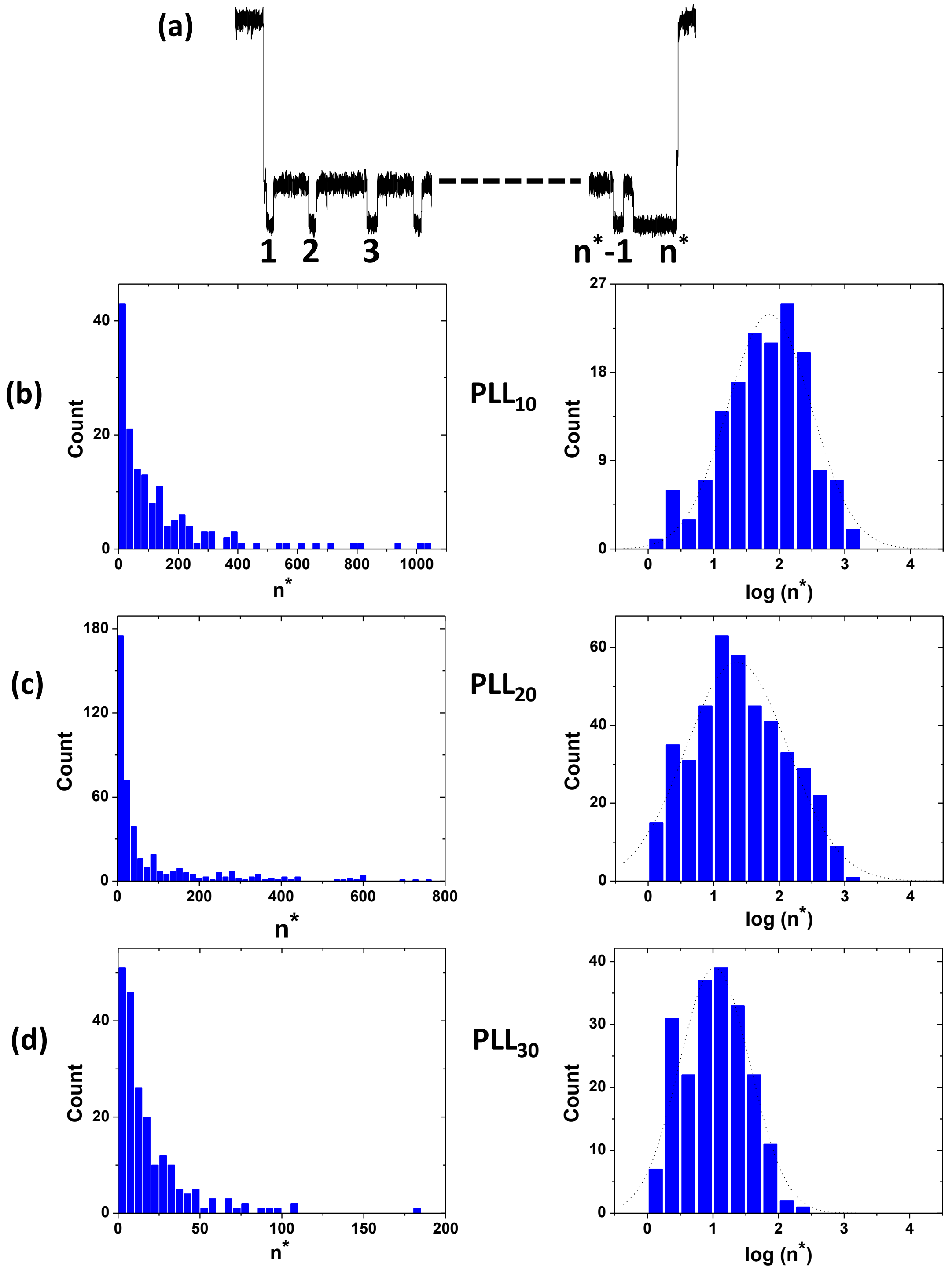


**Figure S26. (a)** Definition of $n^*$ as several binding events in each two-state conductance event. **(b)** Histogram of $n^*$ displaying the number of binding events and Histogram of log $n^*$ for **(b)** $PLL_{10}$ ($n^*_{peak}$ = 71), **(c)** $PLL_{20}$ ($n^*_{peak}$ = 22), and **(d)** $PLL_{30}$ ($n^*_{peak}$ = 10) respectively, the black curve is a log-normal fit to the histogram. The peak n* decreases with PLL length.

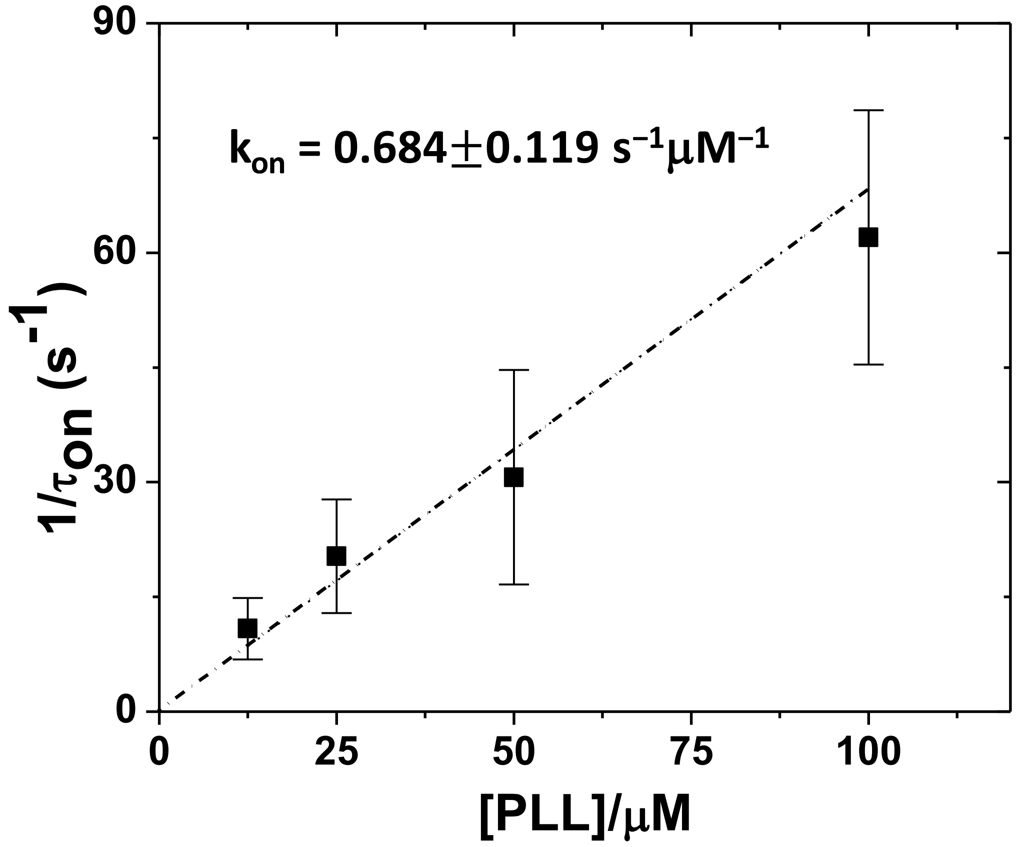


**Figure S27.** The rate constant $k_{on}$ at 140 mV for the association between ssDNA and PLL is obtained from the slope of the linear fit to $1/\tau_{on}$ vs [PLL]. Error bars represents standard deviation obtained from three independent measurements.

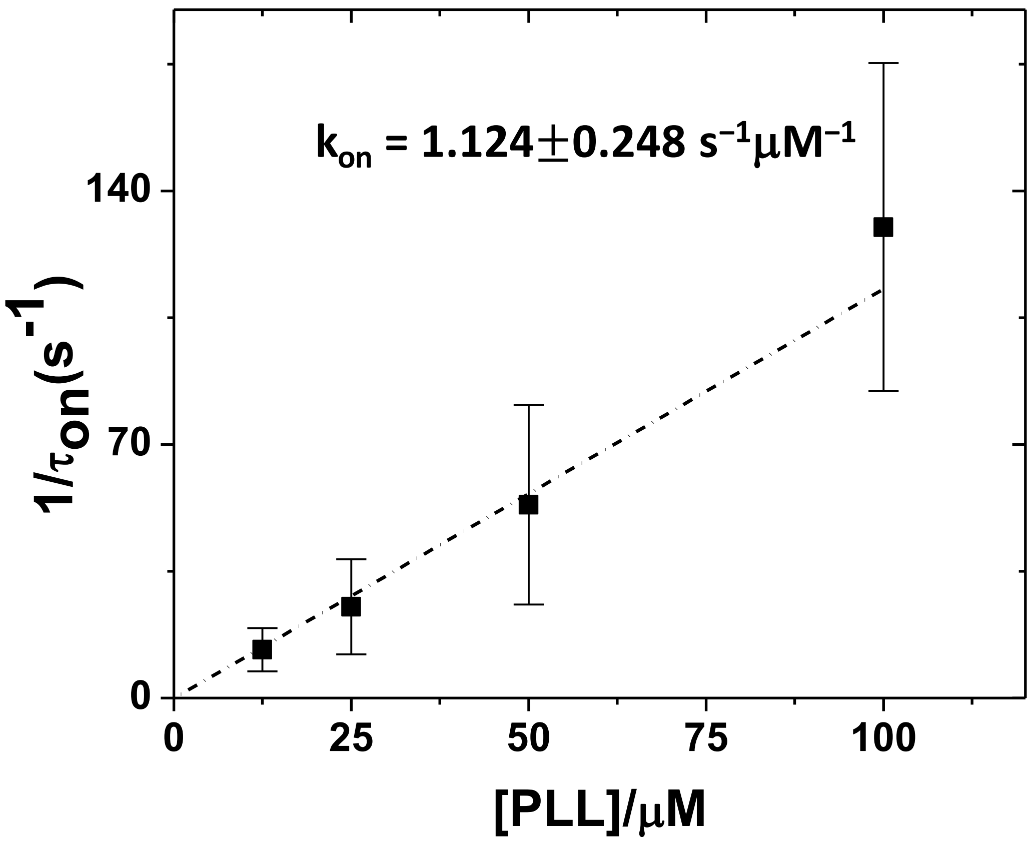


**Figure S28.** The rate constant $k_{on}$ at 160 mV for the association between ssDNA and $PLL_{20}$ is obtained from the slope of the linear fit to $1/\tau_{on}$ vs [PLL]. Error bars represents standard deviation obtained from three independent measurements.

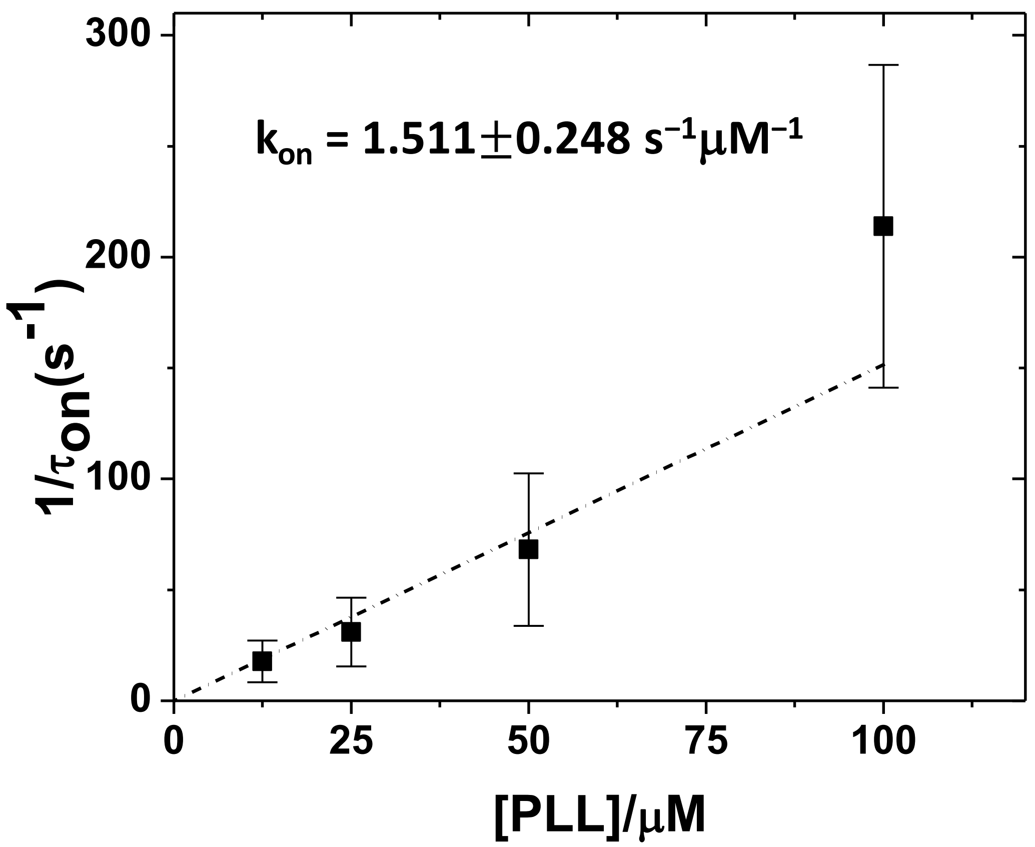


**Figure S29.** The rate constant $k_{on}$ at 180 mV for the association between ssDNA and $PLL_{20}$ is obtained from the slope of the linear fit to $1/\tau_{on}$ vs [PLL]. Error bars represents standard deviation obtained from three independent measurements.

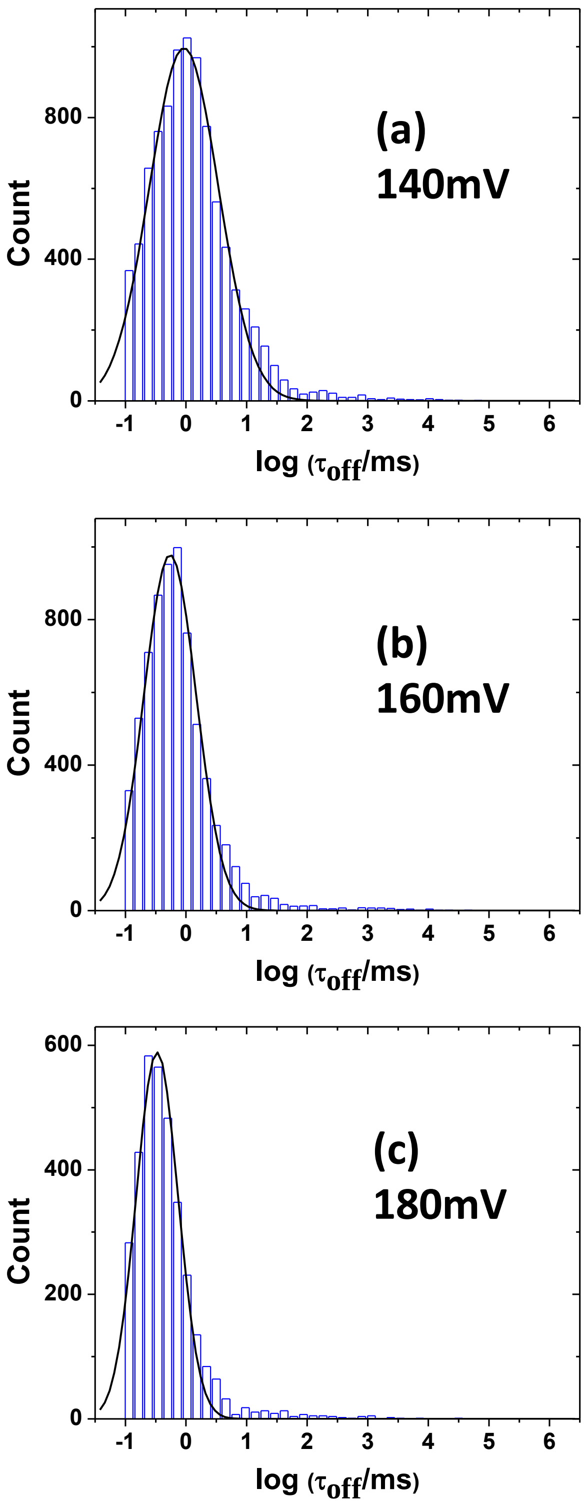


**Figure S30.** Histogram of dwell time of "off" state for $PLL_{20}$ at **(a)** 140 mV **(b)** 160mV, and **(c)** 180mV. The black curve is a log-normal fit to the histogram.

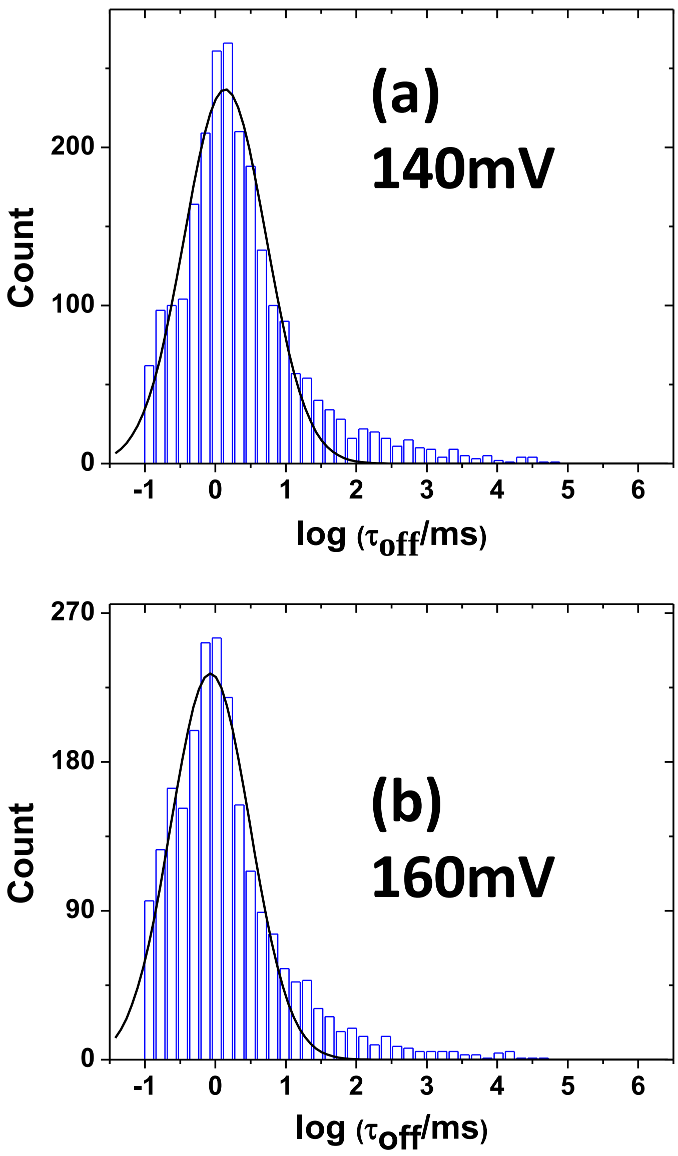


**Figure S31.** Histogram of dwell time of "off" state for $PLL_{30}$ at **(a)** 140 mV **(b)** 160mV. The black curve is a log-normal fit to the histogram.
